# Metagenomics revealing molecular profiling of microbial community structure and metabolic capacity in Bamucuo, Tibet

**DOI:** 10.1101/2022.01.18.476867

**Authors:** Cai Wei, Dan Sun, Wenliang Yuan, Lei Li, Chaoxu Dai, Zuozhou Chen, Xiaomin Zeng, Shihang Wang, Yuyang Zhang, Shouwen Jiang, Zhichao Wu, Dong Liu, Linhua Jiang, Sihua Peng

## Abstract

We performed a survey of the microorganisms in Bamucuo, Tibet, resulting in 160,212 (soil) and 135,994 (water) contigs by shotgun metagenomic methods. We discovered 74 new bacterial species and reconstructed their draft genomes, which were obtained from the 75 reconstructed almost complete metagenomic assembly genomes (MAG) in the soil and water samples. Proteobacteria and Actinobacteria were found to be the most dominant bacterial phyla, while Euryarchaeota was the most dominant archaeal phylum. To our surprise, *Pandoravirus salinus* was found in the soil microbial community. We concluded that the microorganisms in Bamucuo fix carbon mainly through the 3-hydroxypropionic bi-cycle pathway.

**IMPORTANCE:** The Qinghai-Tibet Plateau (QTP) is the highest plateau in the world, and the microorganisms there play vital ecological roles in the global biogeochemical cycle; however, detailed information on the microbial communities in QTP is still lacking, especially in high altitude areas above 4500 meters. This study, for the first time, characterized the microbial community composition and metabolic capacity in QTP high-altitude areas (with an altitude of 4,555 meters), confirmed that QTP is a huge and valuable resource bank in which more new non-resistant antibiotics and many other bioactive substances could be developed. In addition, the discovery of *Pandoravirus salinus* in the soil provides important information for further exploring this unique microorganism, and many draft genomes and the genome annotation information obtained in this study have laid the foundation for further in-depth study of the microbial ecology in Qinghai-Tibet Plateau.

## BACKGROUND

The Qinghai-Tibet Plateau (QTP) is the highest plateau on the earth and is known as the “third Pole” of the world, which is one of the most important water resources in East Asia and plays a vital role in regulating global climate change (1, 2). There are many microorganisms in QTP, which are one of the most important life forms in the extreme environments. Through various metabolic pathways, the microorganisms form the basis of carbon, nitrogen and various nutrient cycles (3, 4), and affect the climate change to a certain extent, such as the emission of the greenhouse gases (CO_2_, CH_4_, N_2_O) (5–7). However, up to now, the diversity information of the microbial communities in QTP are still lacking. To address this issue, we initiated this research by using metagenomics to increase the understanding of the microorganisms in the environments.

With the development of sequencing technology, it is now possible to obtain information from unculturable microorganisms through metagenomics (8). In recent years, the diversity of the microbial communities of the permafrost in QTP has become the focus of most research due to its unique and fragile characteristics (9–12). Han et al. revealed the structure and diversity of Keke Salt Lake microbial community (13). Xing’s research on the soil microbial community in Qaidam Basin showed that the environmental factors are the main driving force for the formation of bacterial community structure (14). However, in QTP with an altitude of more than 4,000 meters, there are few environmental microbiology studies, which are mainly based on 16S Ribosomal RNA Gene Sequencing (16S) rather than shotgun metagenomic sequencing (9, 11–15). Compared with shotgun metagenomic sequencing, 16S has obvious deficiencies: 1) it cannot fully detect the abundance of the microbial community; 2) it can only assign microorganisms to the genus level, but not to specific species; and 3) it only sequence some fragments of the certain regions of the bacterial genome, and the whole microbial genome information cannot be obtained (16, 17).

Shotgun metagenomic sequencing not only provides a stronger and more reliable assessment of microbial diversity, but also provides valuable information on the metabolic potential of microbial communities (18–21). The metagenome-assembled genomes (MAGs) reconstructed from the datasets can help to explore the ecological environment of the QTP more deeply.

Bamucuo is located in QTP, with an altitude of 4,555 meters. The Bamucuo lake is mainly fed by surface runoff surrounding. Here we used shotgun metagenomics to investigate the microbial community in Bamucuo. To the best of our knowledge, this study is the first shotgun metagenomics study of the environmental microbial community at an altitude of more than 4,000 meters in QTP.

## RESULTS

### Quality controlling and metagenome assembling

Shotgun sequencing produced 29,623,109,936 bp and 24,284,563,524 bp of unassembled raw bases from the soil (s05-02) and water (s05-03) samples, respectively (Table 1). In total, 194,202,138 (98.99%) and 159,527,496 (99.19%) reads passed the quality control in the soil and water sample dataset, respectively (Table 1).

**TABLE 1.** Summary of the sample information and the metagenomic sequencing results

|  | s05-02 | s05-03 |
| --- | --- | --- |
| Raw reads | 196,179,536 | 160,824,924 |
| Raw bases (bp) | 29,623,109,936 | 24,284,563,524 |
| Clean reads (%) | 194,202,138(98.99%) | 159,527,496 (99.19%) |
| Clean bases (%) | 29,290,018,832(98.88%) | 24,062,066,782(99.08%) |
| Contigs Num | 160,212 | 135,994 |
| Total length (bp) | 356,933,003 | 397,377,554 |
| Max length (bp) | 833,193 | 286,311 |
| N50 (bp) | 2,289 | 3,841 |
| GC content (%) | 60.18 | 51.33 |
| Average coverage (%) | 31.45 | 45.70 |
| Percent of reference bases covered (%) | 99.90 | 99.92 |

After assembling, a total of 160,212 (soil) and 135,994 (water) contigs were generated. For the soil and water samples, the longest contigs were 833,193 bp and 286,311 bp, GC content were 60.18% and 51.33%, and N50 length were 2,289 bp and 3,841 bp, respectively (Table 1).

After Prokka (22) annotation, 351,242 genes were obtained from the soil sample and 390,943 genes from the water sample (Table 2).

**TABLE 2.** Summary of functional gene annotations against different databases in the two samples

| Database | s05-02 | s05-03 |
| --- | --- | --- |
| Predicted genes | 351,242 | 390,943 |
| nr | 232,679 | 294,919 |
| KEGG | 104,704 | 117,467 |
| COG | 211,757 | 222,635 |
| GO | 45,638 | 59,921 |
| CAZy | 52,427 | 51,940 |
| VFDB | 45,976 | 45,547 |
| BacMet | 21,273 | 18,305 |
| PHI | 59,516 | 60,782 |
| CARD | 11 | 12 |

### Microbial community representation

To reveal the ecological diversity of the high altitude areas, we characterized the microbial structure in Bamucuo. The results showed that bacteria accounting for 99.5% were dominant, with archaea accounting for 0.3% and viruses accounting for 0.08% in the soil. The three most abundant phyla in bacteria were Proteobacteria (54.0%), Actinobacteria (18.6%) and Firmicutes (9.8%), and another 29 phyla were identified (Table S1a). For the most abundant phylum, Proteobacteria, 112 families (Table S1b) and 408 genera were identified (Table S1c). Among the Actinobacteria (Table S1 d, e), 46 families and 141 genera were identified. A total of 2,400 bacterial species were identified (Table S2) in the whole dataset, including *Bacillus cereus* (5.1%), *Hymenobacter sedentarius* (4.5%), *Gemmatirosa kalamazoonesis* (3.0%), etc. In the archaea, Euryarchaeota (81.9%) accounted for the largest proportion, followed by Thaumarchaeota (14.7%) and Crenarchaeota (1.7%). Eighteen species of viruses were identified in the soil sample (Table S3a); Surprisingly, *Pandoravirus salinus* (15.3%), a giant virus, was the most abundant, followed by *Bacillus virus 250* (11.5%), *Mycobacterium phage Jobu08* (7.7%), etc.

The microbial structure in the water was further investigated. The bacteria accounted for the majority (99.1%). Only a small portion was archaea (0.4%), and the rest was viruses (0.5%) (Fig. 2). At the phylum level, Proteobacteria (54.4%), Bacteroidetes (17.9%) and Actinobacteria (15.1%) were the most abundant phyla, and another 30 phyla were identified (Table S4a). Among the Proteobacteria, 124 families (Table S4b) and 464 genera were identified (Table S4c). For the Actinobacteria, 46 families (Table S4d) and 150 genera (Table S4e) were identified. In the water sample, 2,714 bacterial species (Table S5) were identified, in which *Yoonia vestfoldensis* (12.2%), *Belliella baltica* (5.3%) and *Limnohabitans sp. 63ED37-2* (3.6%) were dominant. Among the archaea, Euryarchaeota (98.8%) accounted for the largest proportion. In addition, a total of 52 species of viruses were identified (Table S3b), in which the most abundant was *Chrysochromulina ericina virus* (12.7%), followed by *Phaeocystis globosa virus* (8.8%) and *Synechococcus phage S-PM2* (5.9%).

**FIG 1.**
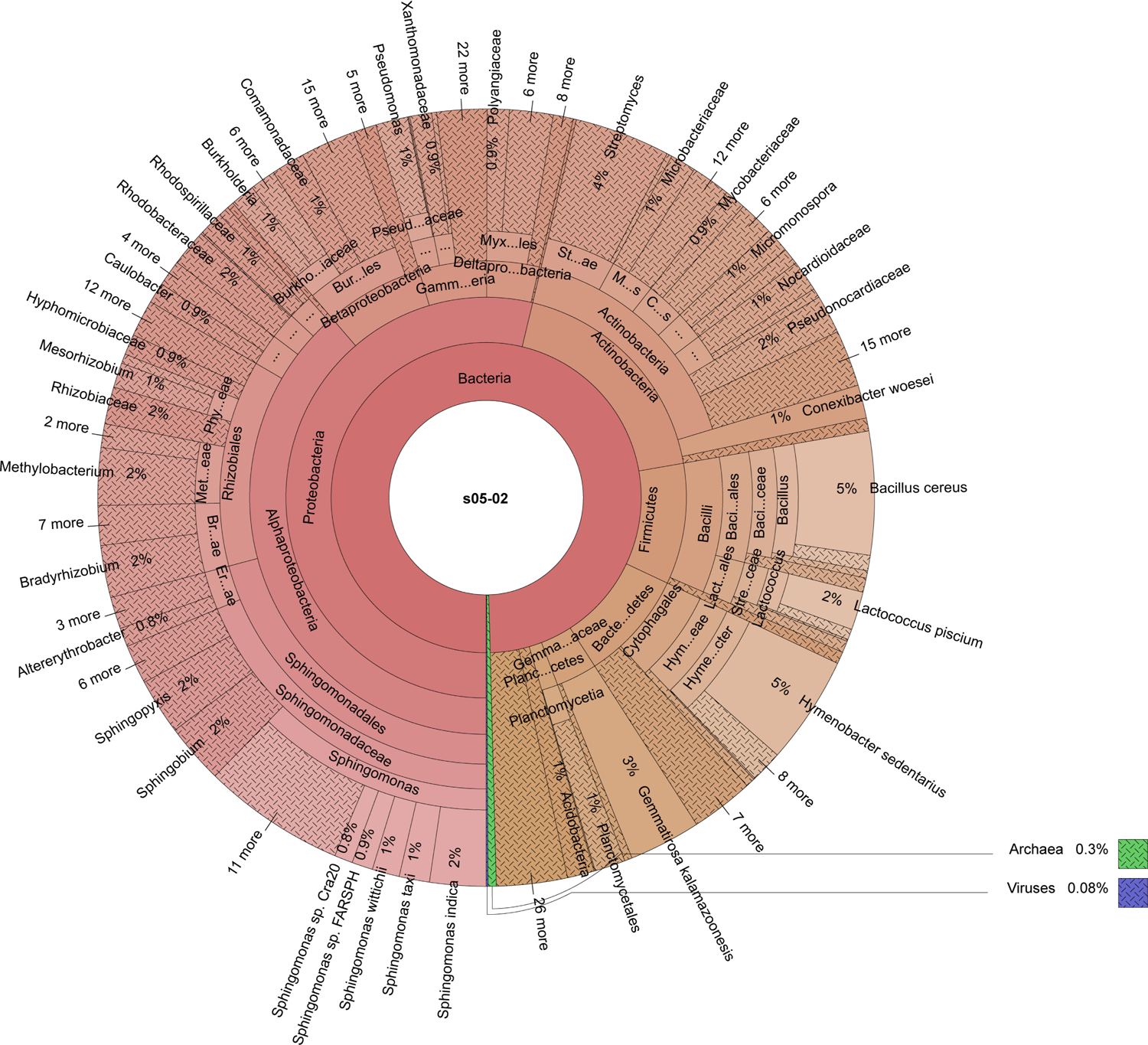
The composition of the soil microbial community. The figure shows the relatively complete distribution at species level.

**FIG 2.**
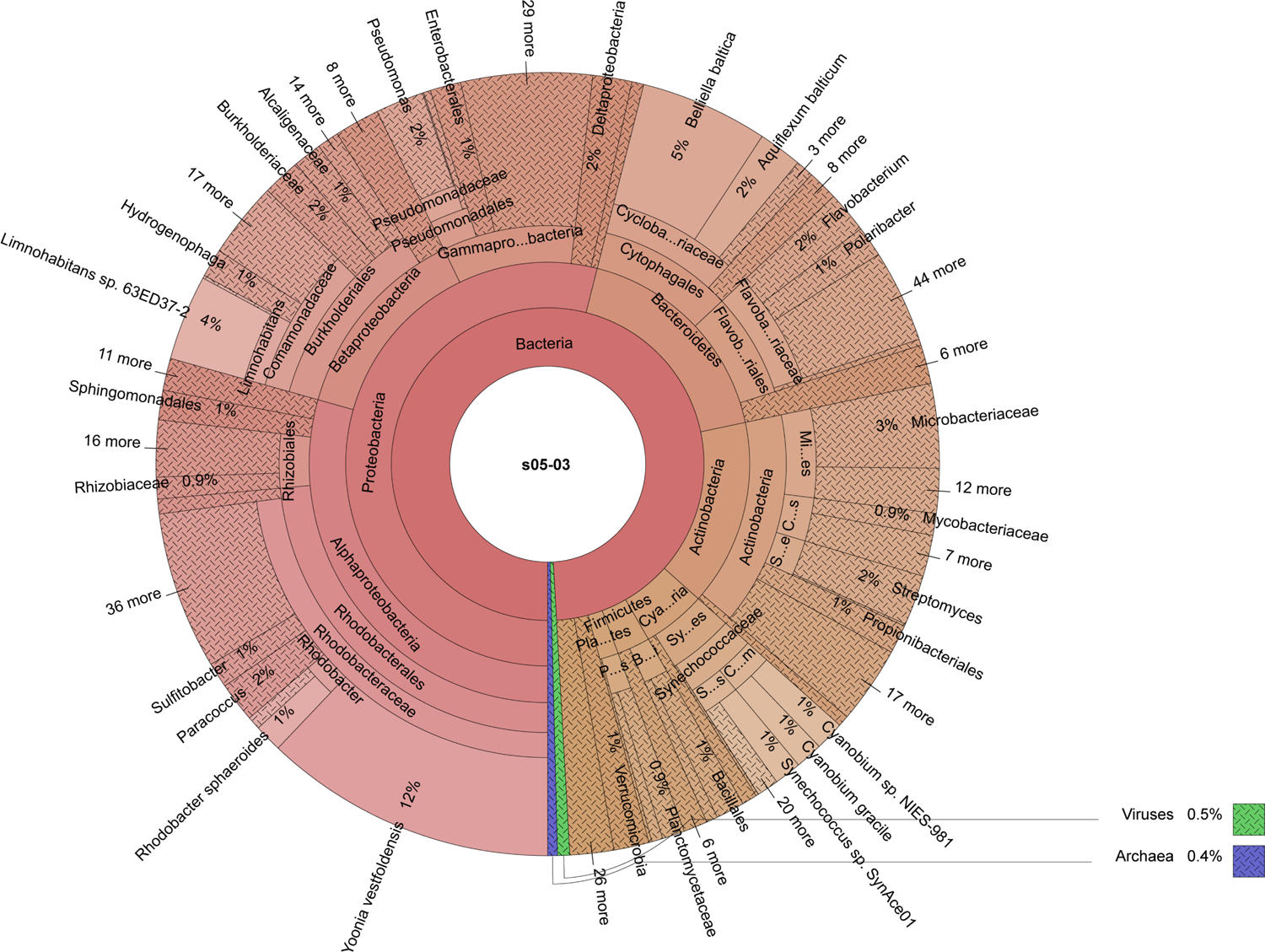
The composition of the water microbial community. The figure shows the most abundant taxa, with the remaining categories to be omitted.

### Environmental microbial function in Bamucuo

We try to characterize the function of the microbial community in Bamucuo. For the soil and water samples, a total of 232,679 and 294,919 genes were annotated against nr database, 104,704 and 117,467 genes against KEGG (23) database, and 45,638 and 59,921 genes against GO (24) database, respectively. More annotation results are shown in Table 2.

The COG (25) functional annotation results are shown in Fig. 3a; Except for function unknown (12.8%), the three most abundant categories in the soil sample were replication, recombination and repair (4.7%), translation, ribosomal structure and biogenesis (3.9%), and transcription (3.9%). In the water sample, except for function unknown (11.0%), translation, ribosomal structure and biogenesis (4.3%) were the most abundant, followed by replication, recombination and repair (3.6%), amino acid transport and metabolism (3.5%), etc.

**FIG 3.**
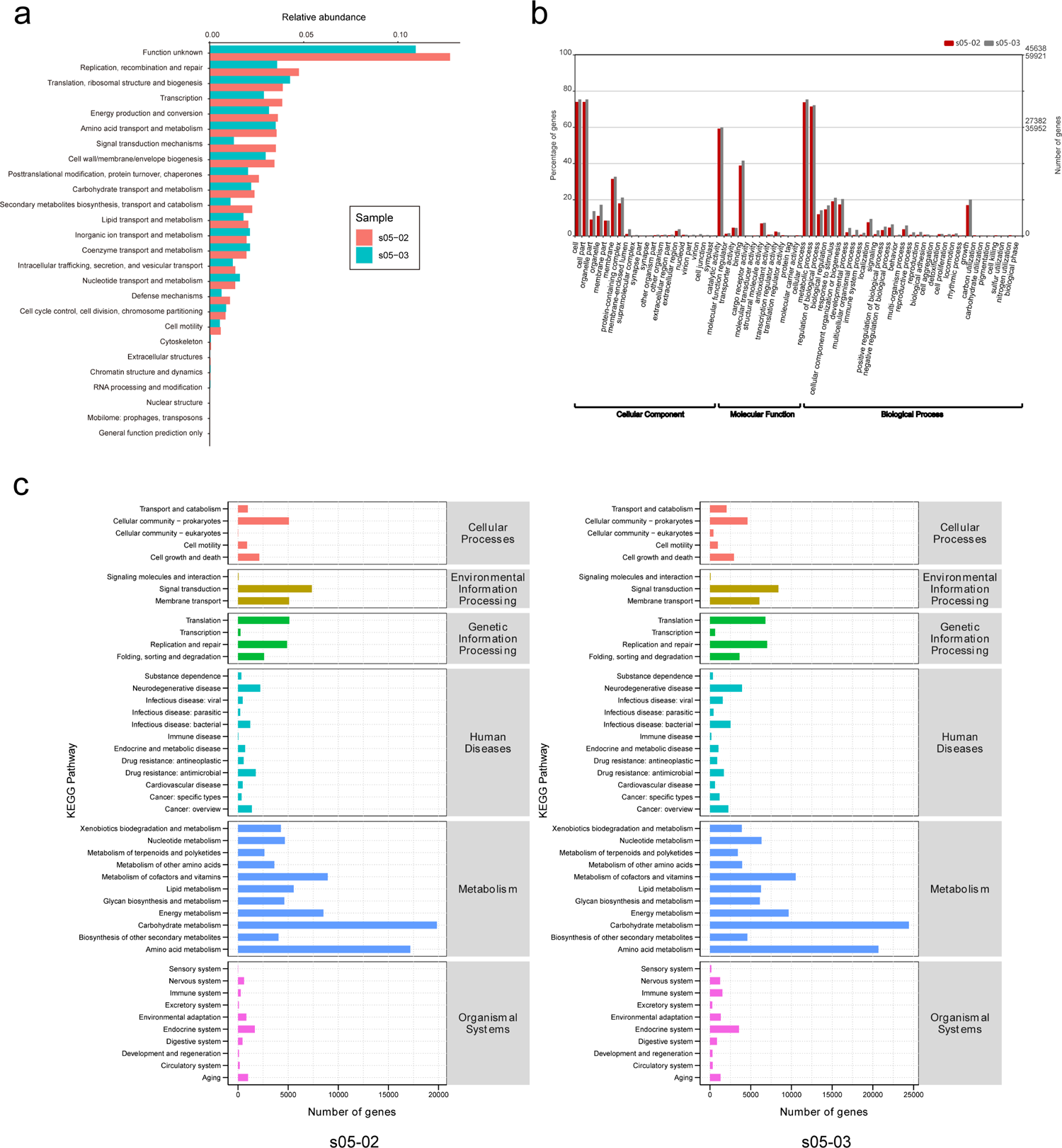
Visualization of functional genes annotation. (a) The relative abundance of the COG (25) functional categories of the two samples. The red bars represent s05-02 (soil), and the green bars represent s05-03 (water). (b) GO distribution analysis of the two samples. The horizontal axis indicates the GO (24) functional classifications, with the red bars representing s05-02 and the grey bars representing s05-03. All the 2-level GO terms are arranged in three main categories (cellular component, molecular function and biological process). (c) The number of the genes annotated at the first two levels KEGG(23) pathway. S05-02 is displayed on the left and s05-03 on the right. The horizontal axes represent the number of the genes, and the vertical axes represent the metabolic pathways.

We investigated the three main categories (cellular component, molecular function and biological process) and the first two levels of the GO terms. In the soil sample, the dominant category was biological process (35.5%), followed by molecular function (33.0%) and cellular component (31.5%) (Fig. 3b). We found that among the biological processes, the top three GO terms were cellular process (33,656), metabolic process (32,600) and response to stimulus (8,678); among the molecular functions, the top three were catalytic activity (27,005), binding (17,692) and structural molecule activity (3,089); and among the cellular components, the top three were cell (33,716), cell part (33,716) and membrane (14,342). Whereas in the water sample, biological process (35.4%) was the dominant category, followed by molecular function (32.7%) and cellular component (31.9%) (Fig. 3b). The top three GO terms were as follows: cellular process (45,113), metabolic process (43,200) and response to stimulus (12,611) among the biological processes; catalytic activity (35,860), binding (24,853) and structural molecule activity (4,247) among the molecular functions; and cell (45,101), cell part (45,101) and membrane (19,524) among the cellular components.

The functional analysis of metagenome showed that for the soil sample, the genes related to metabolism (accounting for 62.2% of the entire gene set annotated) were the most abundant, and the rest were genetic information processing (9.7%), environmental information processing (9.4%), human diseases (7.3%), cellular processes (6.8%), and organismal systems (3.9%). The bar graphs in Fig. 3c represent the number of the annotated genes for each metabolic pathway. Further analysis on the metabolism showed that the most abundant was carbohydrate metabolism (23.6%), followed by amino acid metabolism (20.5%), metabolism of cofactors and vitamins (10.7%), energy metabolism (10.1%), etc. Similarly, for the water sample (Fig. 3c), the most abundant metabolic function module was metabolism (58.4%), followed by human diseases (9.7%), environmental information processing (8.5%), cellular processes (6.4%), organismal systems (6.3%), and genetic information processing (6.2%). In metabolism, carbohydrate metabolism (24.5%) was the most abundant, followed by amino acid metabolism (20.7%), metabolism of cofactors and vitamins (10.6%), energy metabolism (9.7%), etc.

### Reconstruction of the Metagenome-assembled genomes (MAGs)

Metagenomic binning produced 75 reconstructed MAGs with the completeness > 70% and contamination < 10%, including 18 from the soil sample and 57 from the water sample (Fig. 4a and Fig. 4b). A total of 20 MAGs met the high-quality standard with the completeness > 95% and contamination < 5%, accounting for 26.67% of all the MAGs. Table S6 lists the detailed information for the 75 MAGs, including the completeness, contamination, strain heterogeneity, GC content, N50, genomic size and GTDB (26, 27) species classification. The heterogeneity of each MAG was shown in Fig. 4c and Fig. 4d. The original reconstructed MAGs have been reassembled by using reassemble_bins module in MetaWRAP (28) and their genomic integrities have been significantly improved (Fig. 4c and Fig. 4d). The median genome size of the MAGs from soil was 3.58Mb and the median GC content was 62.4% (Table S6), which were similar to the data of the MAGs from water (median value: 3.72Mb; 60.9%).

**FIG 4.**
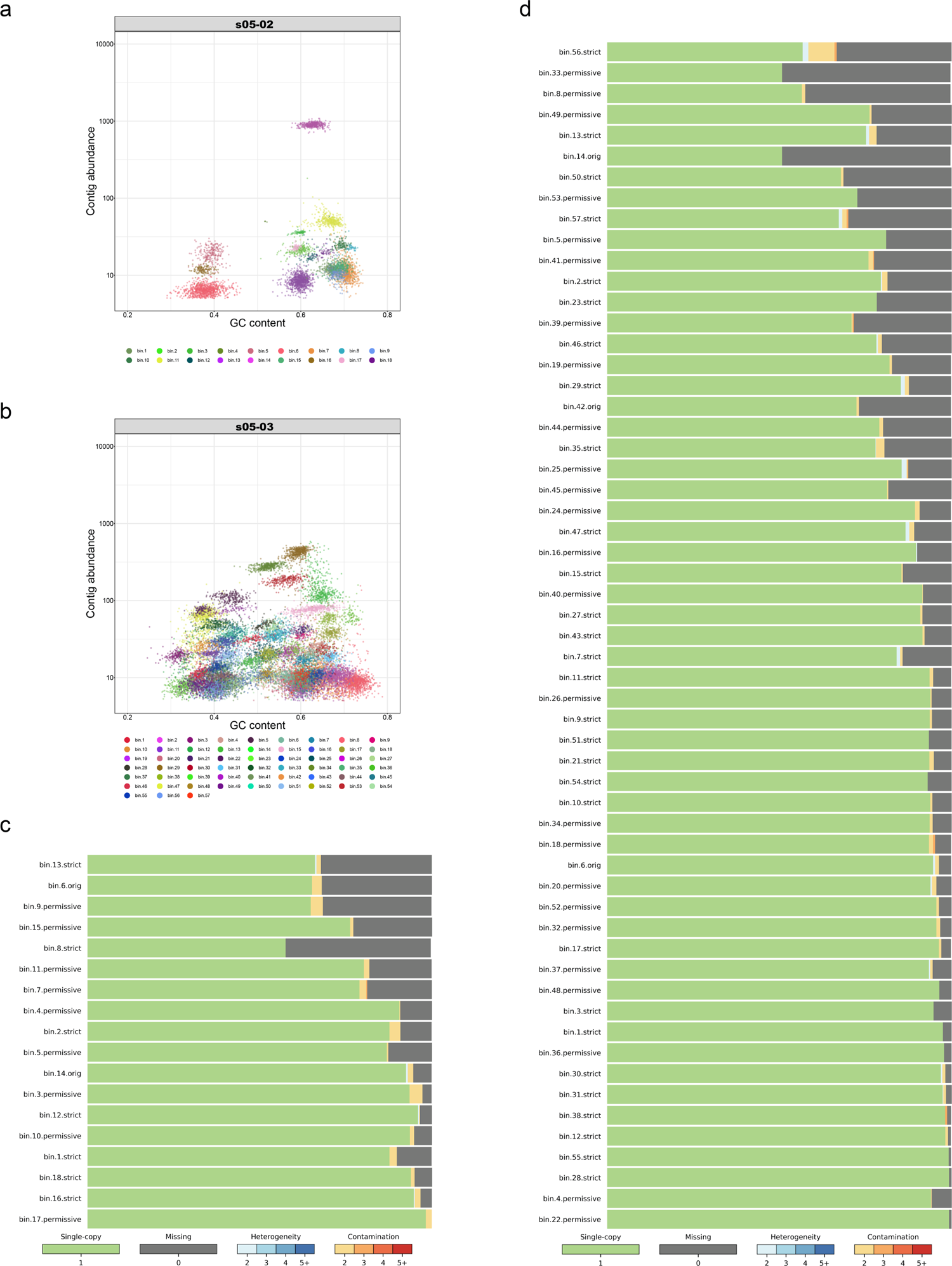
The evaluations of binning and reassembled MAGs. (a) (s05-02, soil) and (b) (s05-03, water): The distribution of the contigs in each bin after refining. The abscissa represents the GC content of the contigs, and the ordinate represents the contig abundance, with one dot representing one contig. The contigs in same color belong to the same bin. (c) (s05-02) and (d) (s05-03): An evaluation of the bins after reassembling using the reassemble_bins module in MetaWRAP (28). The figures indicate Single-copy, Missing, Heterogeneity and Contamination of the reassembled bins.

### Taxonomic identification

To identify the representative microorganisms in Bamucuo, we conducted a comprehensive analysis of each reconstructed MAG. The one-to-one ANI values of all the MAGs were lower than 95%, suggesting that the MAGs were different species. We performed the taxonomic annotation and phylogenetic analysis for the 75 MAGs by using GTDB-tk(29). The phylogenetic tree constructed with the MAGs showed that the 75 MAGs could be taxonomically assigned to 10 phyla (Fig. 5), including Proteobacteria (16 MAGs, five from the soil sample and 11 from the water sample), Bacteroidota (22 MAGs, two from the soil sample and the remaining 20 from the water sample), Actinobacteriota (17 MAGs, three from the soil sample and 14 from the water sample), etc. Searching the GTDB database, the annotation result showed that 74 MAGs were not assigned species names, except for one MAG (the bin 5), which was from the soil sample, with the 98.06% average nucleotide identity (ANI) and 94% alignment fraction (AF), and which was classified as *Lactococcus piscium_C* and highly related to the known *Lactococcus_A piscium_C* (GCF_000981525.1). Obviously, these 74 unspecified MAGs were the draft genomes of the bacterial species newly discovered in our study.

**FIG 5.**
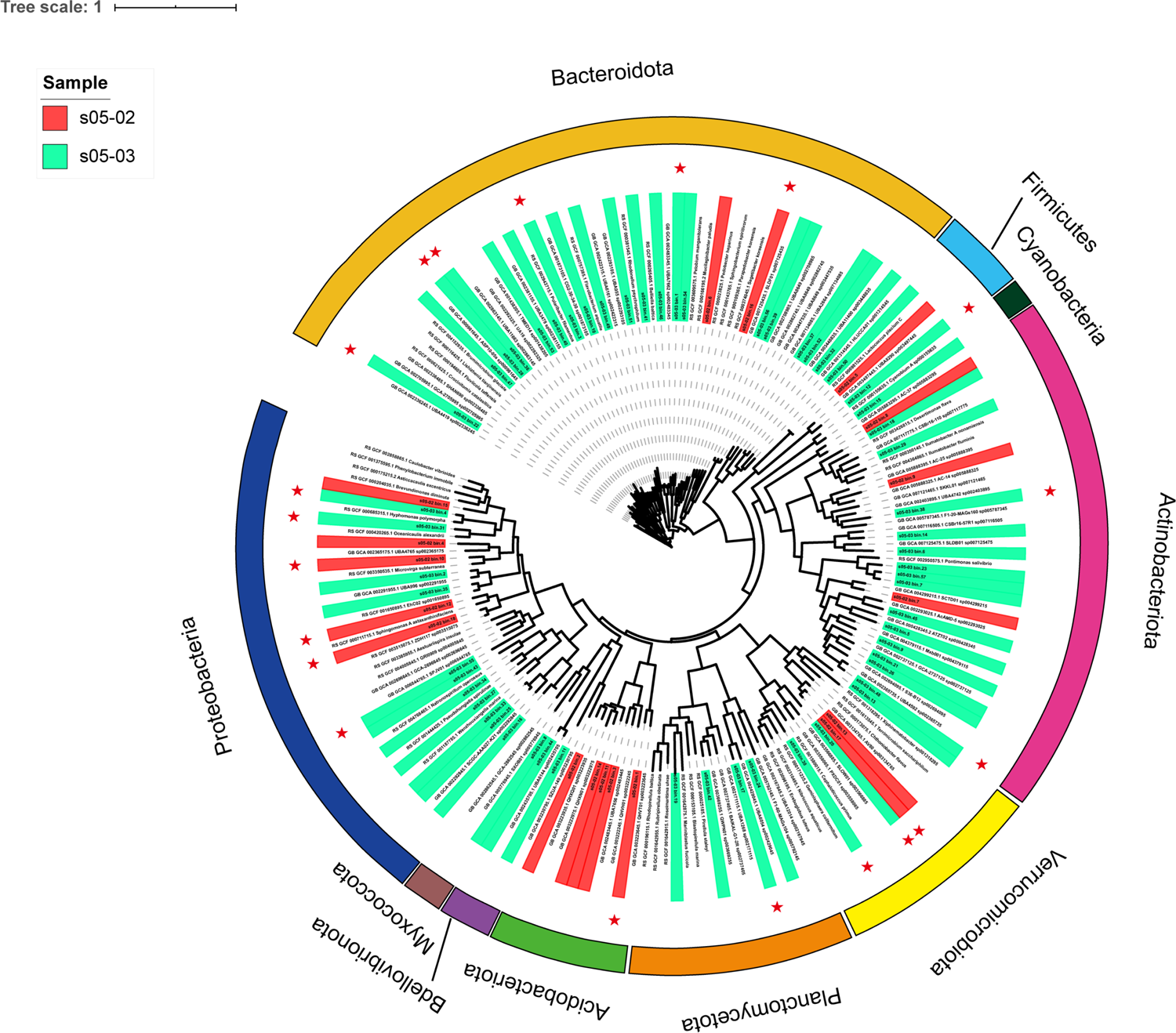
Phylogenetic tree of 75 reconstructed metagenome-assembled genomes (MAGs). The maximum likelihood evolutionary tree is reconstructed for all the 75 MAGs and their nearest neighbor reference genomes. The MAGs of s05-02 were represented by the red background, and the MAGs of s05-03 by the green background. High quality MAGs, namely MAGs with completeness > 95% and contamination < 5%, are marked with a star sign, and the phyla to which they belong were distinguished in outer rings in different colors.

### Function analyses based on the MAGs

To understand the functional potential of the representative microorganisms in Bamucuo, we analyzed the carbon metabolism, methane metabolism, nitrogen metabolism and sulfur metabolism of the reconstructed MAGs.

### Carbon degradation

The 75 MAGs contained abundant hydrolases from the 101 glycoside hydrolases (GH) family (Table S7), among which the MAGs containing GH13 accounted for 77.3% (58 out of 75), and the MAGs containing GH3 and GH23 accounted for 76.0% (57 out of 75).

The enzymes of interest included the starch-degrading enzymes (alpha-amylase, beta-amylase, glucoamylase, alpha-glucosidase, isoamylase and pullulanase), the hemicelluloses-degrading enzymes (xylanase, beta-mannanase, alpha-L-arabinofuranosidase, alpha-D-glucuronidase and beta-xylosidase), the cellulose-degrading enzymes (endo-1,4-glucanase, exo-β-1,4-glucanases and β-1, 4-glucosidases), and the other carbohydrate degrading enzymes.

### Carbon fixation

Almost the 75 MAGs contained the genes encoding the enzymes involved in the 3-hydroxypropionate bi-cycle, reductive tricarboxylic acid cycle, and calvin cycle pathways (Fig.6). The related enzymes are listed in Table S8. A total of 27 MAGs (not listed) contained one or more genes encoding the key enzymes involved in the reductive tricarboxylic acid cycle pathway, e.g., 2-oxoglutarate:ferredoxin oxidoreductase (OGOR), pyruvate:ferredoxin oxidoreductase (PFOR), and ATP-citrate lyase. The genes encoding the key enzyme which is in the calvin cycle pathway, ribulose-1,5-bisphosphate carboxylase-oxygenase (RuBisCO), was only identified in the two MAGs (bin9 in the soil sample and bin32 in the water sample), and another gene encoding the key enzyme involved in this pathway, phosphoribulokinase (PRK), was identified in the two MAGs (bin15 and bin44 in the water sample). In addition, except for the two MAGs (bin9 in the soil sample and bin15 in the water sample), all other MAGs contained the genes encoding the enzymes involved in the reductive acetyl-CoA pathway, and most of the MAGs contained the gene encoding the key enzyme: acetyl-CoA synthase. Except for bin12 in the water sample, 74 MAGs contained the genes related to the dicarboxylate/4-hydroxybutyrate cycle pathway, while 64 MAGs contained the genes encoding the enzymes related to the 3-hydroxypropionate/4-hydroxybutylate cycle pathway.

**FIG 6.**
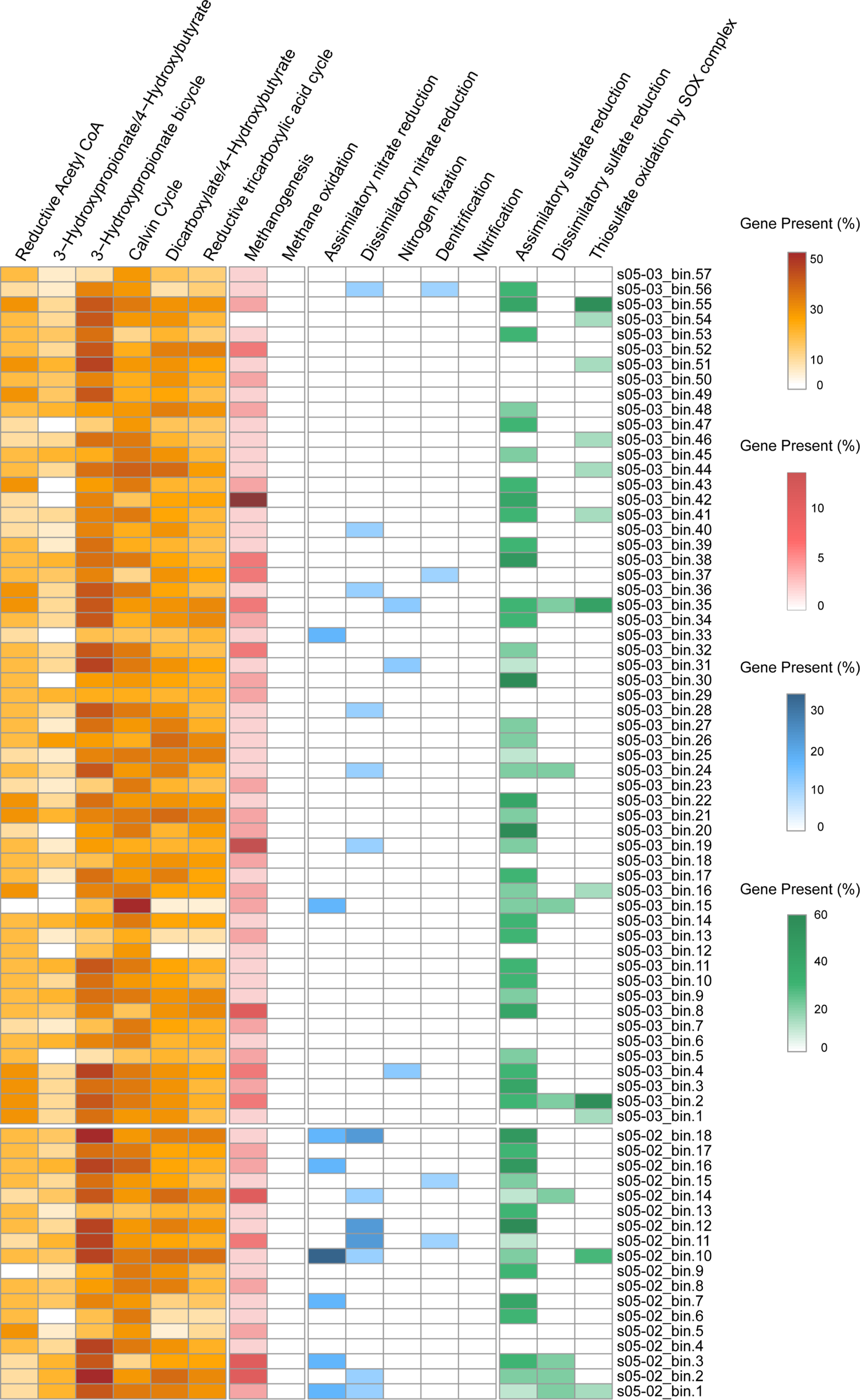
KEGG (23) annotation of energy metabolism for the two samples. The heatmap shows the percentage of the associated genes. The orange boxes represent carbon fixation; the red boxes represent methane metabolism; the blue boxes represent nitrogen metabolism; and the green boxes represent sulfur metabolism. Here, we investigated six carbon fixation pathways, the methanogenesis and methane oxidation pathways, nitrogen metabolism pathways, and sulfur metabolism pathways.

### Methane metabolism

As we know, microbial methanogenesis and methane oxidation play essential roles in the biogeochemical cycles of the earth system. We found that, except bin54 in the water sample, almost all the MAGs contained genes related to the methanogenesis pathway. Fig. 6 shows that the gene methyl coenzyme M reductase (MCR), encoding the key enzyme in the methanogenesis pathway, was found in 17 MAGs (not listed). Furthermore, no gene encoding the enzyme related to the methane oxidation pathway was identified in all the MAGs in the two samples (Fig. 6).

### Nitrogen metabolism

The genes related to the assimilation nitrate reduction pathway were identified in eight MAGs, while 13 MAGs contained the genes encoding the enzymes related to the dissimilation nitrate reduction pathway (Fig. 6). Only one MAG contained the gene encoding respiratory nitrate reductase, a key enzyme, involved in the dissimilation nitrate reduction pathway. Besides, the genes related to the nitrogen fixation pathway were identified in three MAGs (Fig. 6), while the nitrogenase genes (*nif*, *vnf* and *anf*) were not found. Four MAGs (Fig. 6) contained the genes related to the denitrification pathway, while no MAG contained the genes related to the nitrification pathway. Only two MAGs were found to contain the genes encoding nitrite reductase, a key enzyme involved in the denitrification pathway.

### Sulfur metabolism

For sulfur metabolism, 52 MAGs contained the genes related to the assimilatory sulfate reduction pathway; eight MAGs contained the genes related to the dissimilatory sulfate reduction pathway; and 12 MAGs contained the genes related to the thiosulfate oxidation by SOX complex pathway (Fig. 6). Among these 12 MAGs, the marker gene *soxB* related to SOX complex pathway was found in the ten MAGs (bin1 in the soil sample and bin1, bin2, bin35, bin41, bin44, bin46, bin51, bin54, bin55 in the water sample).

### Heavy metal resistance gene identification

To reveal the heavy metal resistance of microorganisms in Bamucuo, we identified common heavy metal resistance genes. The annotation for heavy metal resistance genes revealed diverse resistance genes conferring resistance to 19 heavy metals such as silver, copper, iron, nickel, zinc, and antimony, including *actP*, *copA*, *copB*, *silP*, *acn*, *ctpC*, *ziaA*, etc (Table S8). The results showed that more than 1,064 heavy metal resistance genes were identified in all the MAGs (Fig. 7), among which the heavy metal resistance genes conferring resistance to multi-metals were abundant. Furthermore, the genes conferring resistance to antimony, arsenic, molybdenum and tungsten were identified in all the MAGs.

**FIG 7.**
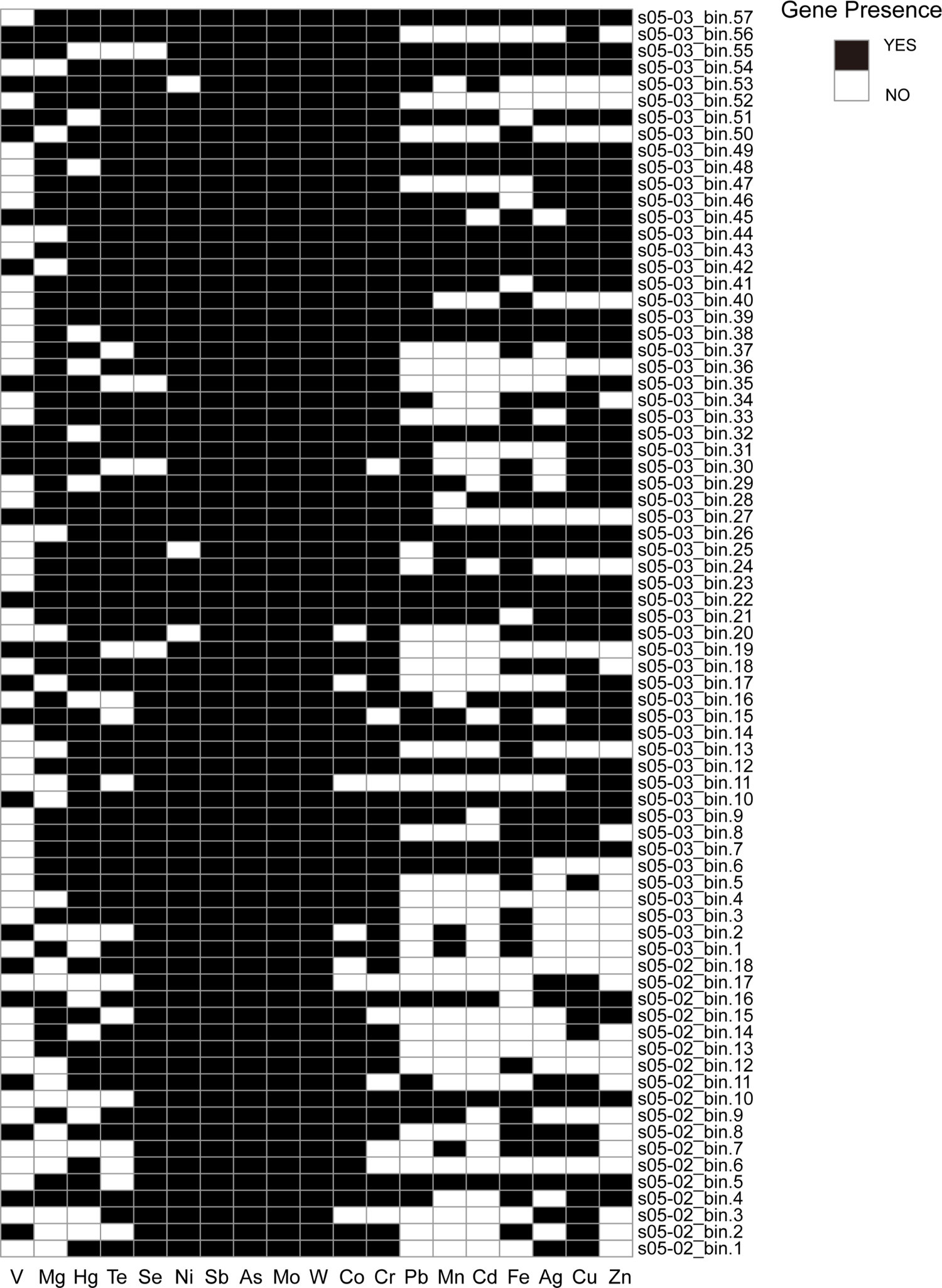
Heavy metal resistance genes identified in the two samples. The black squares represent the presence of the related genes, and the blank squares represent the absence of the related genes. V: the resistant genes for vanadium; Mg: the resistant genes for magnesium; Hg: the resistant genes for mercury; Te: the resistant genes for tellurium; Se: the resistant genes for selenium; Ni: the resistant genes for nickel; Sb: the resistant genes for antimony; As: the resistant genes for arsenic; Mo: the resistant genes for molybdenum; W: the resistant genes for wolfram; Co: the resistant genes for cobalt; Cr: the resistant genes for chromium; Pb: the resistant genes for lead; Mn: the resistant genes for manganese; Cd: the resistant genes for cadmium; Fe: the resistant genes for iron; Ag: the resistant genes for silver; Cu: the resistant genes for copper; Zn: the resistant genes for zinc.

### Biosynthetic gene cluster (BGC) identification

We performed gene cluster identification and annotation for secondary metabolites in all the MAGs. A total of 270 putative BGCs clusters of secondary metabolites were obtained, and 25 types of BGCs were identified in the 75 MAGs (Table S9). Of the 40 NRPSs, 25 existed in the Acidobacteriota MAGs. Terpenes were found in the 61 MAGs, T3PKSs in 17 MAGs, NRPSs in the Acidobacteriota MAGs and Bacteroidota MAGs, PKSs in the Bacteroidota MAGs and Actinobacteria MAGs, and eleven bacteriocins in the Cyanobacteria MAG.

### Antibiotics resistance gene (ARG) identification

To investigate the distribution of ARGs in high altitude areas, we performed ARG analysis based on the MAGs in the two samples. The results showed that the ARGs were found in 15 of the 75 MAGs, including *adeF*, *rpsL* (mycobacterium tuberculosis *rpsL* mutations conferring resistance to streptomycin), etc. Among these 15 MAGs, *adeF* existed in all the soil sample MAGs, and *ampS* existed in bin10. In the water sample, except for *adeF* and *ampS*, *rpsL* accounted for the majority (5 out of 8). The detailed ARGs annotation information is shown in Fig. 8.

**FIG 8.**
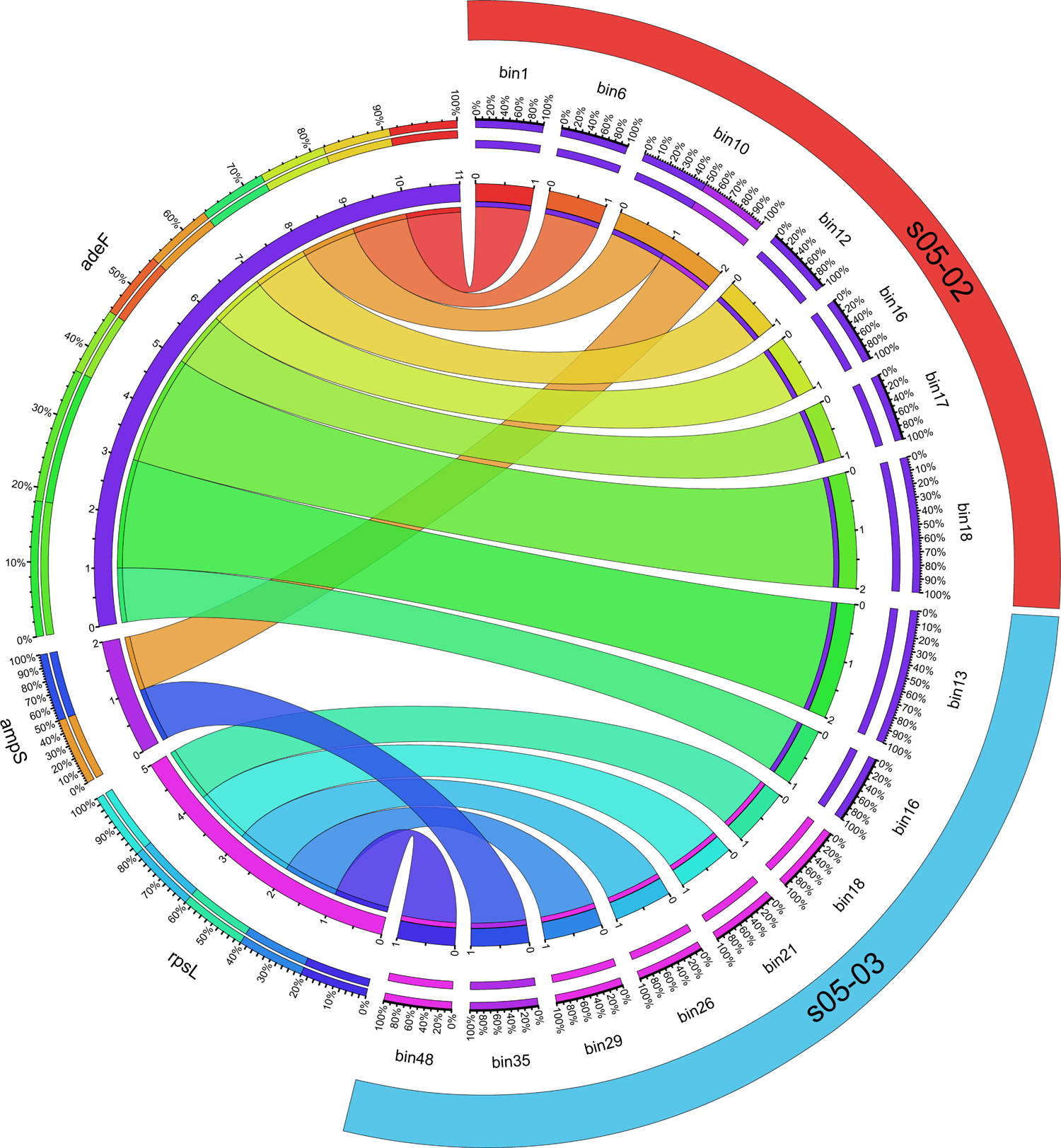
The circos of the annotation of MAGs by using CARD(30). The ARGs information is on the left side of the peripheral ring, and the MAGs information is on the right side. The inner ring shows different MAGs and ARGs by different colors, and the scale represents the abundance information. The width of the band is proportional to the abundance of a particular ARG in each MAG.

## DISCUSSION

### Microbial community structure

The QTP is the highest plateau on the earth, and microbial communities in such extreme environments play an important role in microbial ecology. We have studied the relationship between the distribution and function of the microbial community in Bamcuo at an altitude of 4,555 meters on the Qinghai-Tibet Plateau, revealing the composition of the microbial community. We discovered 74 novel bacterial species and reconstructed 75 MAGs, including 20 high quality MAGs (completeness> 95% and contamination <5%). The median genome size and GC content of the reconstructed MAGs were 2.60Mb and 56.5%, respectively. However, there were still many reads that were not assigned to any bins due to the existence of the sequence data of fungi, eukaryotes, and viruses, which adversely affected our reconstruction of the bacterial MAGs (31).

The dominant bacterial phyla of the microbial communities in our studies were Proteobacteria and Actinobacteria (Fig. 1, Fig. 2 and Fig. 5), which are similar to that in Qaidam Basin in QTP (14) and Qinghai Lake (32). Proteobacteria plays an essential role in various energy metabolism in QTP (14, 33). Actinobacteria is the main source of new antibiotics and bioactive molecular candidates (34). However, many of the existing antibiotics used by human have developed drug resistance (35). Therefore, the discovery of many Actinobacteria in the water and soil microbial communities in Bamucuo may become potential natural resources to develop new antibiotics.

As for archaea, Euryarchaeota was the most abundant in the Bamucuo, which is similar to the observation in Lonar soda Lake in India (36), suggesting that the phylum Euryarchaeota could adapt to the most extreme environmental. However, Thaumarchaeota and Crenarchaeota were only identified from the soil sample in Bamucuo, indicating that there were differences between the soil and water microbial communities. Pandora virus is the largest known virus reported by Philip et al., which was isolated from sediments in the estuary of central Chile and freshwater ponds near Melbourne (Australia) (37), however, in our research, a type of Pandoravirus, Pandoravirus salinus was found in the nearshore soil of Bamucuo. Our finding expanded the habitat of Pandora virus to 4,555 meters above sea level.

### Carbon metabolism

Microorganisms play very important roles in carbon fixation (38). Most genes that play a role in the carbon fixation pathway were found in MAGs (Fig. 6), suggesting that the microbial community in Bamucuo obtains energy through carbon fixation by autotrophic bacteria. Although many different carbon fixation pathways were found in both the samples, the genes related to the 3-hydroxypropionate bi-cycle (9.5%-52.4%), calvin cycle (11.8%-52.9%) and reductive tricarboxylic acid (2.3%-37.2%) pathways were the most complete (Fig. 6), and among these three pathway, the most represented carbon fixation pathway in the 75 MAGs was the 3-hydroxypropionate bi-cycle pathway, which is generally rarely found in most environments because of its high energy cost (39). The abundant genes related to the 3-hydroxypropionate bi-cycle pathway were found in all the 75 MAGs, and the gene encoding the key enzymes, acetyl-CoA carboxylase and propionyl-CoA carboxylase, related to this pathway were present in most MAGs, suggesting that the 3-hydroxypropionate bi-cycle pathway may be dominant under extreme environmental conditions, such as low oxygen, low pressure, low temperature and high salinity. The calvin cycle pathway is the main CO_2_ fixation pathway, which is proved to be widely present in green plants and many autotrophic bacteria in most environments, such as trench (40). However, even if most genes related to the calvin cycle pathway were detected, the genes encoding the key enzymes were not detected in our results. Compared with the calvin cycle pathway, the reductive tricarboxylic acid pathway has lower energy consumption, which was found in certain autotrophic mesophilic bacteria and archaea (39). Our results showed that 27 MAGs contained the genes encoding the key enzymes related to the reductive tricarboxylic acid pathway, indicating that the microorganisms in Bamucuo may prefer to use this pathway instead of the calvin cycle pathway. In addition to the most common three carbon fixation pathways mentioned above, the three less common pathways were the reductive acetyl-CoA (0-30.0%), dicarboxylate/4-hydroxybutyrate (0-39.1%), and 3-hydroxypropionate/4-hydroxybutylate cycle (0-27.8%) pathways. The reductive acetyl-CoA pathway, which requires strict anoxic conditions, plays a role in both psychrophiles and hyperthermophiles(39).

At least 73 MAGs contained the genes encoding the acetyl-CoA synthase, which is a key enzyme related to the reductive acetyl-CoA pathway (Fig. 6), whereas few genes were detected to which were related this pathway. The reductive acetyl-CoA pathway was found to be the most abundant carbon fixation pathway in deep biosphere (41, 42), however, our results suggested that this pathway does not work for the microorganisms in Bamucuo. The 3-hydroxypropionate/4-hydroxybutyrate cycle and the dicarboxylate/4-hydroxybutyrate cycle pathways existed in thermophilic archaea, however, Ruiz-Fernandez et al. concluded that the 3-hydroxypropionate/4-hydroxybutylate cycle is the dominant pathway in anoxic marine zones (43). Obviously, their conclusion is different from ours.

Therefore, the 3-hydroxypropionate bi-cycle pathway is the preferred pathway for microorganisms to fix inorganic carbon into organic carbon in Bamucuo.

### Methane metabolism

In methane metabolism, CO_2_ is converted to CH_4_, which is the second largest greenhouse gas, accounting for about 17% of global warming impact (44), while CO_2_ accounting for about 60% (45). The results showed that almost no genes for the methanogenesis pathway were detected, and the methane oxidation pathway was also not detected (Fig. 6). In the methanogenesis pathway of microbial communities, the methane was generated through the key enzyme, methyl-coenzyme M reductase (MCR), which was identified both in the KEGG (23)annotation results of the MAGs and the metagenomic data, whereas the study on Pangong Lake reported the absence of *MCR* (46). In the contigs of the two samples, the existence of methanogens, such as *Methanobacterium*, *Methanosarcina* and *Methanocaldococcus*, suggested that methanogens can survive in the hypoxic environment in Bamucuo and are crucial in the methane production.

Methane oxidation is an important process to slow down CH_4_ emission (47), which is performed by CH_4_ oxidizing bacteria (methanotrophs). We found that methanotrophs, such as *Methylomonas*, *Methylomicrobium*, and *Methylococcus*, existed in the two samples. However, in the KEGG annotation results, the key functional genes for methane oxidation were not found, which may be due to insufficient sequencing depth.

### Nitrogen cycle

Because the annotation results of the MAGs indicated an incomplete nitrogen metabolism pathway, the nitrogen metabolism might rarely occur in energy metabolism in Bamucuo. Nitrogen loss in QTP is mainly related to denitification (48). In our study, few genes related to the denitrification pathway were identified. However, denitrifying bacteria, such as *Pseudomonas* and *Bacillus*, were identified in both the samples. Nitrification is a key process in biogeochemical nitrogen cycling, which oxidizes ammonia to nitrate via nitrite (49). In our result, nitrosifyer (such as *Nitrosomonas* and *Nitrosococcus*) and nitrifying bacteria (such as *Nitrobacter* and *Nitrpspina*) were low abundant, and the nitrification pathway was not identified, suggesting that nitrification in the surface water and soil microbial communities was relatively rare. However, Rasigraf et al. reported that nitrification and denitrification are the main processes in the Bothnian Sea sediment (50), which is different from our results. It was previously reported that *Proteobacteria* plays an important role in the nitrogen cycle (51), and our results also confirm that the nitrogen fixation pathway has only been found in the *Proteobacteria* MAGs. In addition, the assimilatory nitrate reduction pathway and the dissimilatory nitrate reduction pathway were more abundant than other pathways in the nitrogen cycle, indicating that these two pathways are relatively common in the microbial communities in Bamucuo, which is similar to the results reported by Rathour et al. (46).

### Sulfur cycle

The microbial sulfur cycle has been proposed as the driving force for bacterial survival (52). The sulfur-reducing bacteria (e.g., *Desulfobacterales*, *Desulfovibrionales*, *Desulfotomaculum* and *Desulfosporosinus*) in Bamucuo, which can perform anaerobic respiration utilizing sulfate (SO_4_^2–^) as the terminal electron acceptor for reducing sulfate to hydrogen sulfide (H_2_S), were found in the two samples; similar results in Pangong lake were reported by Rathour et al. (46). The assimilative sulfate reduction pathway was originally reported in plants, however it has been shown to exist in certain bacteria, such as *Allochromatium vinosum* (53). Our results showed that the assimilatory sulfate reduction pathway was relatively complete (Fig. 6) in Bamucuo, while the key functional gene encoding dissimilatory sulfite reductase for dissimilatory sulfate reduction had not been identified, indicating the lack of microorganisms that perform the functions of dissimilatory sulfate reduction.

The sulfur-oxidizing bacteria such as *Sulfuritalea* and *Thiobacillus* were present with a low abundance, and few genes related to the thiosulfate oxidation by SOX complex pathway were identified (Fig. 6). However, the key gene *soxB* related to the thiosulfate oxidation by SOX complex pathway was found in most of the MAGs, so we believed that sulfur oxidation is important in the energy metabolism of the microorganisms in Bamucuo.

### Metal resistance

The microbial community is the main driving force for metal resistance gene distribution, and a previous study showed that these genes are found mainly in the environments contaminated with heavy metals (54). However, in our study, abundant heavy metal resistance genes were identified (Fig. 7). A study supported that the pollution of heavy metals in QTP has been on the rise in recent years (55). In addition, rainwater and runoff may cause microorganisms to accumulate in the lake, being more conducive to the enrichment of heavy metal resistance genes in the water and nearshore soil microbial communities. Chen et al. reported that heavy metals can induce DNA damage and oxidative stress (56). Our study showed that the DNA repair genes (not listed), such as *recB*, *recC*, *recD*, and *recF*, were found in most of the MAGs, which is obviously due to the adaptation of the Bamcuo microbial community to the heavy metal environment.

### BGCs and ARGs

Among the secondary metabolites of microorganisms, special attention was always paid to antibiotics (57). In recent years, some pathogens have developed “drug resistance” to certain antibiotic drugs, and the number of new antibiotics has steadily decreased (35). Therefore, there is an urgent need to find new resources to obtain new antibiotic drugs. Obviously, QTP is the appropriate candidate because of less human activity. Many BGCs were identified in many MAGs (Table S9), suggesting that there may be many novel natural product biosynthetic gene clusters in Bamucuo.

Many terpenoids have important physiological activities and are important sources for the research of natural products and the development of new drugs, and terpenoids are widely distributed in the genomes of many plants, fungi and bacteria(58). In our study, the content of terpenes was the highest (35.56%) (Table S9), however, until now, little is known about their ecological functions in the microbial communities in Bamucuo. Except for terpenes, Non-ribosomal peptide synthases (NRPSs) (14.81%) and polyketide synthases (PKSs) (15.93%) were the most abundant types of BGCs in our study (Table S9), and NRPSs and PKSs are two families of modular mega-synthases (59), which produce compounds such as antibiotics, antifungals, immunosuppressants, and iron-chelating molecules (60). Many NRPS and PKS gene clusters were found in the *Actinobacteria* MAGs, *Acidobacteria* MAGs and *Bacteroidetes* MAGs, suggesting that *Actinobacteria*, *Acidobacteria* and *Bacteroidetes* are the potential natural product biosynthetic producers in Bamucuo. In summary, QTP might be a potential resource bank for discovering new secondary metabolite clusters.

ARG is a new pollutant with mobility (61), and our results showed that few ARGs were identified (Fig. 8) in the soil and water microbial communities in Bamucuo. However, studies in other regions reported that abundant ARGs have been found (36, 62). Therefore, we inferred that Bamucuo has not yet been extensively contaminated by the ARGs. Our study is of great significance in developing new antibiotics without resistance. Interestingly, *rpsL* was only identified in the *Actinobacteriata* MAGs (Fig. 8), indicating that *Actinobacteriata* may be the source of *rpsL* in Bamucuo. Related research also supports that some ARGs found in pathogenic bacteria come from *Actinobacteriata* that produce antibiotics (63).

## CONCLUSIONS

In this study, we investigated the microbial ecosystem of Bamucuo in QTP. The results showed that the water and soil in Bamucuo have unique microbial community structure and metabolic process. Of the 75 MAGs we reconstructed, 74 represented newly discovered bacterial species for the first time, indicating a novel contribution to the global microbial diversity. The presence of *Pandoravirus salinus* in the soil provides important information for further exploring this unique microorganism and discovering more new giant virus. We concluded that the 3-hydroxypropionate bi-cycle pathway is the most representative carbon fixation pathway in Bamucuo. Since *Actinobacteria* was one of the dominant bacteria, QTP can become a valuable resource bank for developing new and non-resistant antibiotic drugs and other bioactive substances. This study provides many draft genomes and the genome annotation information for further in-depth study of the microbial ecology in QTP.

Our future work includes isolation, cultivation, and identification of the 74 new bacterial species discovered in this study. In addition, we will try to obtain the entire genome of the *Pandoravirus salinus* through more in-depth sequencing technology.

## MATERIALS AND METHODS

The detailed sequencing data analysis process of this study is shown in Fig. 9.

**FIG 9.**
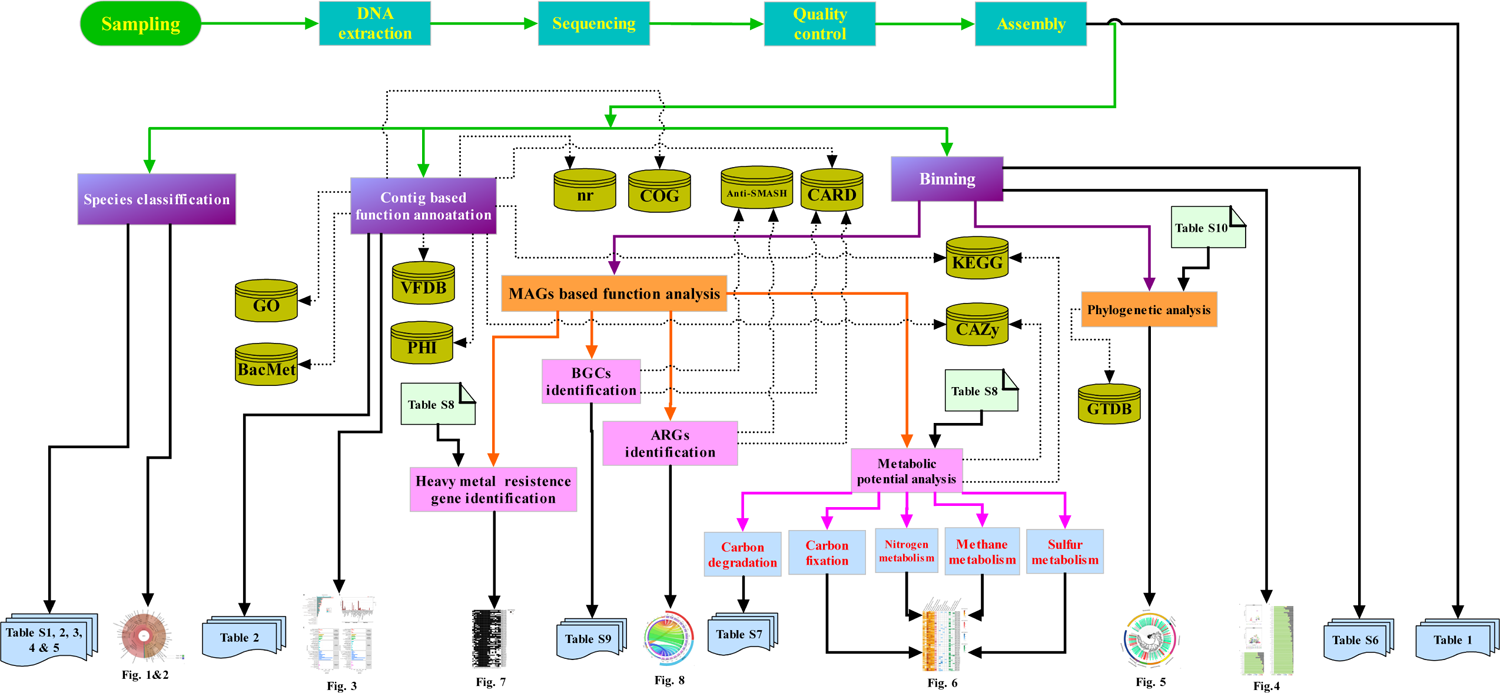
Analysis flow chart in this study. MAGs: metagenome-assembled genomes; BGC: Biosynthetic gene cluster; ARG: Antibiotics resistance gene. The detailed information of the 11 databases in the figure can be referred to the reference list.

### Sampling site and sample collection

The soil and water samples involved in this study were collected on June 20, 2017 in Bamucuo (31°34′N, 90°6′E), at an altitude of 4,555 meters, Bango County, Nagqu Prefecture, Tibet, China. We collected 30L water sample at depth of 0.5m through a filter membrane with a pore diameter of 0.22μm, and the water temperature was 13℃ when sampling. The ambient temperature was 19°C. The soil sample was collected from 0.2m below the surface, and then stored in ice boxes for bringing back to the laboratory. For simplicity, in this article, we refer to these two samples as s05-02 (soil) and s05-03 (water).

### DNA extraction and sequencing

According to the manufacturer’s instructions, genomic DNA was extracted using OMEGA’s DNA extraction kit (http://www.omegabiotek.com.cn/, Guangzhou, China) and some modifications were made in the experimental procedure. The purity and concentration of the extracted genomic DNA were measured on NanoDrop 2000 spectrophotometer (Thermo Fisher Scientific, Massachusetts, United States), and the quality of the extracted genomic DNA was detected by 1% agarose gel electrophoresis. The total genomic DNA was stored in a −20 ℃ refrigerator. Then, the genomic DNA was sent to Shanghai Meiji Biomedical Technology Co., Ltd for sequencing. The length of the library was 300bp, and Hiseq 4000 high-throughput sequencer was used for sequencing.

### Metagenomic assembly and gene abundance analysis

The raw reads were processed using fastp v. 0.19.5 (64) for removing adapter sequences at the 3’ and 5’ terminal. The reads with the length less than 50 bp after shearing, the reads with an average quality value (Phred value) less than 20, and the reads containing N bases were all removed. FastQC v. 0.11.9(65) was used to perform the sequence quality control. The assembly module of MetaWRAP v. 1.2.1(28) was then used for assembling, which include the assembly software MEGAHIT v 1.1.3 (66), setting the parameters as follows: minimum contig length of 1000 bp, and k-mer sizes of 21, 29, 39, 59, 79, 99, 119 and 141, respectively. BBMap v. 38.87 (https://sourceforge.net/projects/bbmap/) was used to map clean reads to each contig for calculating the coverage information. Prokka v. 1.14.6 (22) was used to annotate the assembled sequence, with all the parameters to be the default values. Salmon v. 1.3.0 (67) was then used to estimate the genetic abundance for each sample.

### Species classification and functional annotation

We integrated the annotation results obtained by Kraken2 v. 2.1.1 (68) and Bracken v. 2.6.0 (69), viewing the results of the species annotation by Krona v. 2.7.1 (70).

Using Diamond v. 2.0.5(71) as a aligner, the alignments (e-value≤1E-5) were performed against Non-Redundant Protein Sequence Database (NCBI-nr), Carbohydrate-Active Enzyme Database (CAZy) (72), Pathogen-Host Interactions Database (PHI) (73), Virulence Factors Database (VFDB) (74), and Biocide and Metal Resistance Genes Database (BacMet) (75), respectively. Rgi v. 5.1.1, based on Comprehensive Antibiotic Resistance Database (CARD) (30), was employed to predict the resistomes. The Kyoto Encyclopedia of Genes and Genomes (KEGG) (23) annotation was performed using KAAS online analysis website (76), and ggplot2 was used to display the annotation at the first two levels of KEGG (23). Clusters of Orthologous Groups of Proteins (COG) (25) and Gene Ontology (GO) (24, 77, 78) functional annotation were performed using emapper v. 2.0.1(79) based on the eggNOG orthology data (80), and then Web Gene Ontology Annotation Plot (WEGO v. 2.0) (81) was used to classify the GO functions.

### Metagenomic binning

Three modules (MetaBAT2(82), MaxBin2 (83), and CONCOCT (84)), included in MetaWRAP (28), were used to perform the binning operation for all the clean reads, setting the minimum contig length to be 1500, and then three different binning results were obtained. CheckM v 1.0.12 (85) was employed to calculate the completeness and contamination (42, 86). Then, bin_refinement module in MetaWRAP was used to obtain the bins with the completeness of more than 70% and with the contamination of less than 10% (parameter: -c 70 -x 10), and these bins were called metagenome-assembled genomes (MAGs).

By using the MetaWRAP pipeline (28), the clean reads of the each sample were mapped to the assembled MAGs by using the quant_bins module; the relative abundance of all the MAGs was estimated; a preliminary gene annotation for each MAG was obtained by using the annotate_bins module; and the distribution of the contigs in each bin was visualized by the blobology module.

### Taxonomic classification and phylogenetic analysis

fastANI v. 1.32 (87) was used to cluster the MAGs, and the MAGs with an ANI value less than 95% were considered to be different species of bacteria. GTDB-tk v. 1.3.0 (29) “classify” workflow was used to conduct species classification and to calculate the ANI values of the MAGs based on Genome Taxonomy Database (GTDB) (26, 27). For phylogenetic analysis, we first searched the GTDB database to find the corresponding reference genomes which are closest to each MAG (Table S10), and then used GTDB-tk (29) “align” method to perform multiple sequence alignments for the marker genes in the MAGs and the marker genes in the corresponding reference genomes. Importing the above alignment results, the “infer” method (FastTree v. 2.1.10) (88) was used to construct a phylogenetic tree under the Whelan-And-Goldman (WAG) model, with the parameters to be set to “default”. Finally, the annotation information was refined using ITOL v. 5.7 (89).

### Functional analysis of MAGs

The carbon degradation enzymes were identified using dbCAN(90). We examined the KEGG (23) biochemical maps for the six known carbon fixation pathways (reductive acetyl-CoA, 3-hydroxypropionate/4-hydroxybutylate cycle, 3-hydroxypropionate bi-cycle, dicarboxylate/4-hydroxybutyrate cycle, reductive tricarboxylic acid cycle, and calvin cycle (42, 91)) and identified the genes encoding the enzymes involved in methanogenesis and methane oxidation. The genes related to nitrogen and sulfur metabolism were investigated (Table S8). Diamond (71) was used to perform the alignments for all the MAGs (e-value < 1E-5) against BacMet Database (75). The MAGs were submitted to the Anti-SMASH v. 5.2.0 (92, 93) web site to screen the secondary metabolite biosynthetic gene clusters (BGCs) of these draft genomes, with the model set to ‘strict’. Rgi (30) was used to predict the resistomes for the MAGs.

## SUPPLEMENTAL MATERIAL

**Table S1.** Detailed information in the soil microbial community at phylum, family, and genus level

**Table S2.** The information of the bacteria annotated in the soil microbial community at species level

**Table S3.** The information of the viruses identified in Bamucuo at species level

**Table S4.** Detailed information in the water microbial community at phylum, family, and genus level

**Table S5.** The information of the bacteria annotated in the water microbial community at species level

**Table S6.** Table S6. Statistics of the completeness, contamination, GC content, N50, size, and GTDB species classification of all the reconstructed MAGs

**Table S7.** Presence of glycoside hydrolases family assigned to CAZy database in all the MAGs

**Table S8.** Complete list of EC number and genes used for identification of the metabolic pathway and the heavy metal resistome

**Table S9.** The biosynthetic gene clusters (BGCs) in all the MAGs

**Table S10.** List of the selected reference genomes used to construct the phylogenetic tree

## AVAILABILITY OF Data

All the raw sequencing data were deposited in the NCBI Sequence Read Archive (SRA) under accession numbers SAMN17838283 (soil sample-05-02) and SAMN17838284 (water sample-05-03). The metagenomic sequence data is bundled under NCBI BioProject number PRJNA700111.

## ACKNOWLEDGEMENTS

This research was supported by the Shanghai Natural Science Foundation (SNSF, grant no. 15zr1420800) to Sihua Peng, and partly supported by the National Science Foundation (NSF, grant no. 61775139) to Linhua Jiang. S.H.P., D.L., and L.H.J. conceived of the study. C.W., D.S., W.L.Y., and L.L. participated in the study design. C.W., D.S., C.X.D., and S.H.W performed the wet-lab experiments. C.W., D.S., W.L.Y., Z.Z.C and Y.Y.Z. processed the raw data and performed the computational analyses. S.W.J. and Z.C.W designed the sampling scheme and guided the process of DNA extraction and metagenome sequencing. S.H.P and X.M.Z. guided the pipelines of all the analyses. C.W. wrote the first draft of the manuscript. S.H.P., D.L., and L.H.J. reviewed and revised the manuscript. All authors reviewed and approved the final version of the manuscript.

## REFERENCES

1. Chen BX, Zhang XZ, Tao J, Wu JS, Wang JS, Shi P, Zhang YJ, Yu CQ. 2014. The impact of climate change and anthropogenic activities on alpine grassland over the Qinghai-Tibet Plateau. Agricultural and Forest Meteorology 189–190:11-18. https://doi.org/10.1016/j.agrformet.2014.01.002.

2. Kuang XX, Jiao JJ. 2016. Review on climate change on the Tibetan Plateau during the last half century. Journal of Geophysical Research: Atmospheres 121:3979–4007. https://doi.org/10.1002/2015JD024728.

3. Rui JP, Li JB, Wang SP, An JX, Liu WT, Lin QY, Yang YF, He ZL, Li XZ. 2015. Responses of Bacterial Communities to Simulated Climate Changes in Alpine Meadow Soil of the Qinghai-Tibet Plateau. Applied and Environmental Microbiology 81:6070–6077. https://doi.org/10.1128/aem.00557-15.

4. Cavicchioli R, Ripple WJ, Timmis KN, Azam F, Bakken LR, Baylis M, Behrenfeld MJ, Boetius A, Boyd PW, Classen AT, Crowther TW, Danovaro R, Foreman CM, Huisman J, Hutchins DA, Jansson JK, Karl DM, Koskella B, Welch DBM, Martiny JBH, Moran MA, Orphan VJ, Reay DS, Remais JV, Rich VI, Singh BK, Stein LY, Stewart FJ, Sullivan MB, van Oppen MJH, Weaver SC, Webb EA, Webster NS. 2019. Scientists’ warning to humanity: microorganisms and climate change. Nature Reviews Microbiology 17:569–586. https://doi.org/10.1038/s41579-019-0222-5.

5. Raymond PA, Hartmann J, Lauerwald R, Sobek S, McDonald C, Hoover M, Butman D, Striegl R, Mayorga E, Humborg C, Kortelainen P, Durr H, Meybeck M, Ciais P, Guth P. 2013. Global carbon dioxide emissions from inland waters. Nature 503:355–359. https://doi.org/10.1038/nature12760.

6. DelSontro T, Beaulieu JJ, Downing JA. 2018. Greenhouse gas emissions from lakes and impoundments: Upscaling in the face of global change. Limnology and Oceanography Letters 3:64–75. https://doi.org/10.1002/lol2.10073.

7. Saunois M, Bousquet P, Poulter B, Peregon A, Ciais P, Canadell JG, Dlugokencky EJ, Etiope G, Bastviken D, Houweling S, Janssens-Maenhout G, Tubiello FN, Castaldi S, Jackson RB, Alexe M, Arora VK, Beerling DJ, Bergamaschi P, Blake DR, Brailsford G, Brovkin V, Bruhwiler L, Crevoisier C, Crill P, Covey K, Curry C, Frankenberg C, Gedney N, Hoglund-Isaksson L, Ishizawa M, Ito A, Joos F, Kim HS, Kleinen T, Krummel P, Lamarque JF, Langenfelds R, Locatelli R, Machida T, Maksyutov S, McDonald KC, Marshall J, Melton JR, Morino I, Naik V, O’Doherty S, Parmentier FJW, Patra PK, Peng CH, Peng SS, et al. 2016. The global methane budget 2000-2012. Earth System Science Data 8:697–751. https://doi.org/10.5194/essd-8-697-2016.

8. Jousset A, Bienhold C, Chatzinotas A, Gallien L, Gobet A, Kurm V, Kuesel K, Rillig MC, Rivett DW, Salles JF, van der Heijden MGA, Youssef NH, Zhang X, Wei Z, Hol WHG. 2017. Where less may be more: how the rare biosphere pulls ecosystems strings. Isme Journal 11:853–862. https://doi.org/10.1038/ismej.2016.174.

9. Hu WG, Zhang Q, Tian T, Li DY, Cheng G, Mu J, Wu QB, Niu FJ, An LZ, Feng HY. 2016. Characterization of the prokaryotic diversity through a stratigraphic permafrost core profile from the Qinghai-Tibet Plateau. Extremophiles 20:337–349. https://doi.org/10.1007/s00792-016-0825-y.

10. Hu WG, Zhang Q, Li DY, Cheng G, Mu J, Wu QB, Niu FJ, An LZ, Feng HY. 2014. Diversity and community structure of fungi through a permafrost core profile from the Qinghai-Tibet Plateau of China. Journal of Basic Microbiology 54:1331–1341. https://doi.org/10.1002/jobm.201400232.

11. Wei SP, Cui HP, He H, Hu F, Su X, Zhu YH. 2014. Diversity and Distribution of Archaea Community along a Stratigraphic Permafrost Profile from Qinghai-Tibetan Plateau, China. Archaea-an International Microbiological Journal 12:240817. https://doi.org/10.1155/2014/240817.

12. Chen YL, Deng Y, Ding JZ, Hu HW, Xu TL, Li F, Yang GB, Yang YH. 2017. Distinct microbial communities in the active and permafrost layers on the Tibetan Plateau. Molecular Ecology 26:6608–6620. https://doi.org/10.1111/mec.14396.

13. Han R, Zhang X, Liu J, Long QF, Chen LS, Liu DL, Zhu DR. 2017. Microbial community structure and diversity within hypersaline Keke Salt Lake environments. Canadian Journal of Microbiology 63:895–908. https://doi.org/10.1139/cjm-2016-0773.

14. Xing R, Gao QB, Zhang FQ, Wang JL, Chen SL. 2019. Large-scale distribution of bacterial communities in the Qaidam Basin of the Qinghai-Tibet Plateau. Microbiologyopen 8:e909. https://doi.org/10.1002/mbo3.909.

15. Ma AA, Zhang XF, Jiang K, Zhao CM, Liu JL, Wu MD, Wang Y, Wang MM, Li JH, Xu SJ. 2020. Phylogenetic and Physiological Diversity of Cultivable Actinomycetes Isolated From Alpine Habitats on the Qinghai-Tibetan Plateau. Frontiers in Microbiology 11:555351. https://doi.org/10.3389/fmicb.2020.555351.

16. Ranjan R, Rani A, Metwally A, McGee HS, Perkins DL. 2016. Analysis of the microbiome: Advantages of whole genome shotgun versus 16S amplicon sequencing. Biochemical and Biophysical Research Communications 469:967–977. https://doi.org/10.1016/j.bbrc.2015.12.083.

17. Sunagawa S, Mende DR, Zeller G, Izquierdo-Carrasco F, Berger SA, Kultima JR, Coelho LP, Arumugam M, Tap J, Nielsen HB, Rasmussen S, Brunak S, Pedersen O, Guarner F, de Vos WM, Wang J, Li JH, Dore J, Ehrlich SD, Stamatakis A, Bork P. 2013. Metagenomic species profiling using universal phylogenetic marker genes. Nature Methods 10:1196–1199. https://doi.org/10.1038/nmeth.2693.

18. Zhou JZ, He ZL, Yang YF, Deng Y, Tringe SG, Alvarez-Cohen L. 2015. High-Throughput Metagenomic Technologies for Complex Microbial Community Analysis: Open and Closed Formats. Mbio 6:e02288–14. https://doi.org/10.1128/mBio.02288-14.

19. Sharpton TJ. 2014. An introduction to the analysis of shotgun metagenomic data. Frontiers in Plant Science 5:209. https://doi.org/10.3389/fpls.2014.00209.

20. Wiseschart A, Mhuantong W, Tangphatsornruang S, Chantasingh D, Pootanakit K. 2019. Shotgun metagenomic sequencing from Manao-Pee cave, Thailand, reveals insight into the microbial community structure and its metabolic potential. Bmc Microbiology 19:144. https://doi.org/10.1186/s12866-019-1521-8.

21. Alneberg J, Karlsson CMG, Divne AM, Bergin C, Homa F, Lindh MV, Hugerth LW, Ettema TJG, Bertilsson S, Andersson AF, Pinhassi J. 2018. Genomes from uncultivated prokaryotes: a comparison of metagenome-assembled and single-amplified genomes. Microbiome 6:173. https://doi.org/10.1186/s40168-018-0550-0.

22. Seemann T. 2014. Prokka: rapid prokaryotic genome annotation. Bioinformatics 30:2068–2069. https://doi.org/10.1093/bioinformatics/btu153.

23. Kanehisa M, Goto S. 2000. KEGG: Kyoto Encyclopedia of Genes and Genomes. Nucleic Acids Research 28:27–30. https://doi.org/10.1093/nar/28.1.27.

24. Ashburner M, Ball CA, Blake JA, Botstein D, Butler H, Cherry JM, Davis AP, Dolinski K, Dwight SS, Eppig JT, Harris MA, Hill DP, Issel-Tarver L, Kasarskis A, Lewis S, Matese JC, Richardson JE, Ringwald M, Rubin GM, Sherlock G, Gene Ontology C. 2000. Gene Ontology: tool for the unification of biology. Nature Genetics 25:25–29. https://doi.org/10.1038/75556.

25. Tatusov RL, Koonin EV, Lipman DJ. 1997. A genomic perspective on protein families. Science 278:631–637. https://doi.org/10.1126/science.278.5338.631.

26. Parks DH, Chuvochina M, Chaumeil PA, Rinke C, Mussig AJ, Hugenholtz P. 2020. A complete domain-to-species taxonomy for Bacteria and Archaea. Nature Biotechnology 38:1079–1086. https://doi.org/10.1038/s41587-020-0501-8.

27. Parks DH, Chuvochina M, Waite DW, Rinke C, Skarshewski A, Chaumeil PA, Hugenholtz P. 2018. A standardized bacterial taxonomy based on genome phylogeny substantially revises the tree of life. Nature Biotechnology 36:996–1004. https://doi.org/10.1038/nbt.4229.

28. Uritskiy GV, DiRuggiero J, Taylor J. 2018. MetaWRAP-a flexible pipeline for genome-resolved metagenomic data analysis. Microbiome 6:158. https://doi.org/10.1186/s40168-018-0541-1.

29. Chaumeil PA, Mussig AJ, Hugenholtz P, Parks DH. 2020. GTDB-Tk: a toolkit to classify genomes with the Genome Taxonomy Database. Bioinformatics 36:1925–1927. https://doi.org/10.1093/bioinformatics/btz848.

30. Alcock BP, Raphenya AR, Lau TTY, Tsang KK, Bouchard M, Edalatmand A, Huynh W, Nguyen ALV, Cheng AA, Liu SH, Min SY, Miroshnichenko A, Tran HK, Werfalli RE, Nasir JA, Oloni M, Speicher DJ, Florescu A, Singh B, Faltyn M, Hernandez-Koutoucheva A, Sharma AN, Bordeleau E, Pawlowski AC, Zubyk HL, Dooley D, Griffiths E, Maguire F, Winsor GL, Beiko RG, Brinkman FSL, Hsiao WWL, Domselaar GV, McArthur AG. 2020. CARD 2020: antibiotic resistome surveillance with the comprehensive antibiotic resistance database. Nucleic Acids Research 48:D517–D525. https://doi.org/10.1093/nar/gkz935.

31. Nayfach S, Roux S, Seshadri R, Udwary D, Varghese N, Schulz F, Wu DY, Paez-Espino D, Chen IM, Huntemann M, Palaniappan K, Ladau J, Mukherjee S, Reddy TBK, Nielsen T, Kirton E, Faria JP, Edirisinghe JN, Henry CS, Jungbluth SP, Chivian D, Dehal P, Wood-Charlson EM, Arkin AP, Tringe SG, Visel A, Woyke T, Mouncey NJ, Ivanova NN, Kyrpides NC, Eloe-Fadrosh EA, Consortium I-MD. 2021. A genomic catalog of Earth’s microbiomes. Nature Biotechnology 39:499–509. https://doi.org/10.1038/s41587-020-0718-6.

32. Huang QY, Briggs BR, Dong HL, Jiang HC, Wu G, Edwardson C, De Vlaminck I, Quake S. 2014. Taxonomic and Functional Diversity Provides Insight into Microbial Pathways and Stress Responses in the Saline Qinghai Lake, China. Plos One 9:e111681. https://doi.org/10.1371/journal.pone.0111681.

33. Wang R, Han R, Long QF, Gao X, Xing JW, Shen GP, Zhu DR. 2020. Bacterial and Archaeal Communities within an Ultraoligotrophic, High-altitude Lake in the Pre-Himalayas of the Qinghai-Tibet Plateau. Indian Journal of Microbiology 60:363–373. https://doi.org/10.1007/s12088-020-00881-8.

34. Genilloud O. 2017. Actinomycetes: still a source of novel antibiotics. Natural Product Reports 34:1203–1232. https://doi.org/10.1039/c7np00026j.

35. Ventola CL. 2015. The antibiotic resistance crisis: part 1: causes and threats. P & T: a peer-reviewed journal for formulary management 40:277–83.

36. Chakraborty J, Sapkale V, Rajput V, Shah M, Kamble S, Dharne M. 2020. Shotgun metagenome guided exploration of anthropogenically driven resistomic hotspots within Lonar soda lake of India. Ecotoxicology and Environmental Safety 194:110443. https://doi.org/10.1016/j.ecoenv.2020.110443.

37. Philippe N, Legendre M, Doutre G, Coute Y, Poirot O, Lescot M, Arslan D, Seltzer V, Bertaux L, Bruley C, Garin J, Claverie JM, Abergel C. 2013. Pandoraviruses: Amoeba Viruses with Genomes Up to 2.5 Mb Reaching That of Parasitic Eukaryotes. Science 341:281–286. https://doi.org/10.1126/science.1239181.

38. La Cono V, Ruggeri G, Azzaro M, Crisafi F, Decembrini F, Denaro R, La Spada G, Maimone G, Monticelli LS, Smedile F, Giuliano L, Yakimov MM. 2018. Contribution of Bicarbonate Assimilation to Carbon Pool Dynamics in the Deep Mediterranean Sea and Cultivation of Actively Nitrifying and CO2-Fixing Bathypelagic Prokaryotic Consortia. Frontiers in Microbiology 9:3. https://doi.org/10.3389/fmicb.2018.00003.

39. Berg IA. 2011. Ecological Aspects of the Distribution of Different Autotrophic CO2 Fixation Pathways. Applied and Environmental Microbiology 77:1925–1936. https://doi.org/10.1128/aem.02473-10.

40. Zhang XX, Xu W, Liu Y, Cai MW, Luo ZH, Li M. 2018. Metagenomics Reveals Microbial Diversity and Metabolic Potentials of Seawater and Surface Sediment From a Hadal Biosphere at the Yap Trench. Frontiers in Microbiology 9:2402. https://doi.org/10.3389/fmicb.2018.02402.

41. Magnabosco C, Ryan K, Lau MCY, Kuloyo O, Lollar BS, Kieft TL, van Heerden E, Onstott TC. 2016. A metagenomic window into carbon metabolism at 3 km depth in Precambrian continental crust. Isme Journal 10:730–741. https://doi.org/10.1038/ismej.2015.150.

42. Momper L, Jungbluth SP, Lee MD, Amend JP. 2017. Energy and carbon metabolisms in a deep terrestrial subsurface fluid microbial community. Isme Journal 11:2319–2333. https://doi.org/10.1038/ismej.2017.94.

43. Ruiz-Fernandez P, Ramirez-Flandes S, Rodriguez-Leon E, Ulloa O. 2020. Autotrophic carbon fixation pathways along the redox gradient in oxygen-depleted oceanic waters. Environmental Microbiology Reports 12:334–341. https://doi.org/10.1111/1758-2229.12837.

44. Cai YF, Zheng Y, Bodelier PLE, Conrad R, Jia ZJ. 2016. Conventional methanotrophs are responsible for atmospheric methane oxidation in paddy soils. Nature Communications 7:11728. https://doi.org/10.1038/ncomms11728.

45. Nakicenovic N, Alcamo J, Grubler A, Riahi K, Roehrl RA, Rogner HH, Victor N. 2000. Special Report on Emissions Scenarios, Working Group III, Intergovernmental Panel on Climate Change. Cambridge University Press.

46. Rathour R, Gupta J, Mishra A, Rajeev AC, Dupont CL, Thakur IS. 2020. A comparative metagenomic study reveals microbial diversity and their role in the biogeochemical cycling of Pangong lake. Science of the Total Environment 731:139074. https://doi.org/10.1016/j.scitotenv.2020.139074.

47. Canelhas MR, Denfeld BA, Weyhenmeyer GA, Bastviken D, Bertilsson S. 2016. Methane oxidation at the water-ice interface of an ice-covered lake. Limnology and Oceanography 61:S78–S90. https://doi.org/10.1002/lno.10288.

48. Wang SY, Liu WY, Zhao SY, Wang C, Zhuang LJ, Liu L, Wang WD, Lu YL, Li FB, Zhu GB. 2019. Denitrification is the main microbial N loss pathway on the Qinghai-Tibet Plateau above an elevation of 5000 m. Science of the Total Environment 696:133852. https://doi.org/10.1016/j.scitotenv.2019.133852.

49. Spasov E, Tsuji JM, Hug LA, Doxey AC, Sauder LA, Parker WJ, Neufeld JD. 2020. High functional diversity among Nitrospira populations that dominate rotating biological contactor microbial communities in a municipal wastewater treatment plant. Isme Journal 14:1857–1872. https://doi.org/10.1038/s41396-020-0650-2.

50. Rasigraf O, Schmitt J, Jetten MSM, Luke C. 2017. Metagenomic potential for and diversity of N-cycle driving microorganisms in the Bothnian Sea sediment. Microbiologyopen 6:e475. https://doi.org/10.1002/mbo3.475.

51. Ren M, Zhang ZF, Wang XL, Zhou ZW, Chen DL, Zeng H, Zhao SM, Chen LL, Hu YL, Zhang CY, Liang YX, She QX, Zhang Y, Peng N. 2018. Diversity and Contributions to Nitrogen Cycling and Carbon Fixation of Soil Salinity Shaped Microbial Communitiesin Tarim Basin. Frontiers in Microbiology 9:431. https://doi.org/10.3389/fmicb.2018.00431.

52. Cao HL, Wang Y, Lee OO, Zeng X, Shao ZZ, Qian PY. 2014. Microbial Sulfur Cycle in Two Hydrothermal Chimneys on the Southwest Indian Ridge. Mbio 5:e00980–13. https://doi.org/10.1128/mBio.00980-13.

53. Neumann S, Wynen A, Truper HG, Dahl C. 2000. Characterization of the cys gene locus from Allochromatium vinosum indicates an unusual sulfate assimilation pathway. Molecular Biology Reports 27:27–33. https://doi.org/10.1023/a:1007058421714.

54. Song WJ, Qi R, Zhao L, Xue NN, Wang LY, Yang YY. 2019. Bacterial community rather than metals shaping metal resistance genes in water, sediment and biofilm in lakes from arid northwestern China. Environmental Pollution 254:113041. https://doi.org/10.1016/j.envpol.2019.113041.

55. Wu J, Lu J, Li LM, Min XY, Luo YM. 2018. Pollution, ecological-health risks, and sources of heavy metals in soil of the northeastern Qinghai-Tibet Plateau. Chemosphere 201:234–242. https://doi.org/10.1016/j.chemosphere.2018.02.122.

56. Chen Y, Jiang YM, Huang HY, Mou LC, Ru JL, Zhao JH, Xiao S. 2018. Long-term and high-concentration heavy-metal contamination strongly influences the microbiome and functional genes in Yellow River sediments. Science of the Total Environment 637:1400–1412. https://doi.org/10.1016/j.scitotenv.2018.05.109.

57. Atanasov AG, Zotchev SB, Dirsch VM, Supuran CT, Int Nat Prod Sci T. 2021. Natural products in drug discovery: advances and opportunities. Nature Reviews Drug Discovery 20:200-216. https://doi.org/10.1038/s41573-020-00114-z.

58. Yamada Y, Kuzuyama T, Komatsu M, Shin-ya K, Omura S, Cane DE, Ikeda H. 2015. Terpene synthases are widely distributed in bacteria. Proceedings of the National Academy of Sciences of the United States of America 112:857–862. https://doi.org/10.1073/pnas.1422108112.

59. Cuadrat RRC, Ionescu D, Davila AMR, Grossart HP. 2018. Recovering Genomics Clusters of Secondary Metabolites from Lakes Using Genome-Resolved Metagenomics. Frontiers in Microbiology 9:251. https://doi.org/10.3389/fmicb.2018.00251.

60. Finking R, Marahiel MA. 2004. Biosynthesis of nonribosomal peptides. Annual Review of Microbiology 58:453–488. https://doi.org/10.1146/annurev.micro.58.030603.123615.

61. Ma YJ, Wilson CA, Novak JT, Riffat R, Aynur S, Murthy S, Prudens A. 2011. Effect of Various Sludge Digestion Conditions on Sulfonamide, Macrolide, and Tetracycline Resistance Genes and Class I Integrons. Environmental Science & Technology 45:7855–7861. https://doi.org/10.1021/es200827t.

62. Pan X, Lin L, Zhang WH, Dong L, Yang YY. 2020. Metagenome sequencing to unveil the resistome in a deep subtropical lake on the Yunnan-Guizhou Plateau, China. Environmental Pollution 263:114470. https://doi.org/10.1016/j.envpol.2020.114470.

63. Jiang XL, Ellabaan MMH, Charusanti P, Munck C, Blin K, Tong YJ, Weber T, Sommer MOA, Lee SY. 2017. Dissemination of antibiotic resistance genes from antibiotic producers to pathogens. Nature Communications 8:15784. https://doi.org/10.1038/ncomms15784.

64. Chen SF, Zhou YQ, Chen YR, Gu J. 2018. fastp: an ultra-fast all-in-one FASTQ preprocessor. Bioinformatics 34:884–890. https://doi.org/10.1093/bioinformatics/bty560.

65. Cock PJA, Fields CJ, Goto N, Heuer ML, Rice PM. 2010. The Sanger FASTQ file format for sequences with quality scores, and the Solexa/Illumina FASTQ variants. Nucleic Acids Research 38:1767–1771. https://doi.org/10.1093/nar/gkp1137.

66. Li DH, Luo RB, Liu CM, Leung CM, Ting HF, Sadakane K, Yamashita H, Lam TW. 2016. MEGAHIT v1.0: A fast and scalable metagenome assembler driven by advanced methodologies and community practices. Methods 102:3–11. https://doi.org/10.1016/j.ymeth.2016.02.020.

67. Patro R, Duggal G, Love MI, Irizarry RA, Kingsford C. 2017. Salmon provides fast and bias-aware quantification of transcript expression. Nature Methods 14:417–419. https://doi.org/10.1038/nmeth.4197.

68. Wood DE, Lu J, Langmead B. 2019. Improved metagenomic analysis with Kraken 2. Genome Biology 20:257. https://doi.org/10.1186/s13059-019-1891-0.

69. Lu J, Breitwieser FP, Thielen P, Salzberg SL. 2017. Bracken: estimating species abundance in metagenomics data. Peerj Computer Science doi:10.7717/peerj-cs.104:e104. https://doi.org/10.7717/peerj-cs.104.

70. Ondov BD, Bergman NH, Phillippy AM. 2011. Interactive metagenomic visualization in a Web browser. Bmc Bioinformatics 12:385. https://doi.org/10.1186/1471-2105-12-385.

71. Buchfink B, Xie C, Huson DH. 2015. Fast and sensitive protein alignment using DIAMOND. Nature Methods 12:59–60. https://doi.org/10.1038/nmeth.3176.

72. Lombard V, Ramulu HG, Drula E, Coutinho PM, Henrissat B. 2014. The carbohydrate-active enzymes database (CAZy) in 2013. Nucleic Acids Research 42:D490–D495. https://doi.org/10.1093/nar/gkt1178.

73. Winnenburg R, Baldwin TK, Urban M, Rawlings C, Kohler J, Hammond-Kosack KE. 2006. PHI-base: a new database for pathogen host interactions. Nucleic Acids Research 34:D459–D464. https://doi.org/10.1093/nar/gkj047.

74. Chen LH, Zheng DD, Liu B, Yang J, Jin Q. 2016. VFDB 2016: hierarchical and refined dataset for big data analysis-10 years on. Nucleic Acids Research 44:D694–D697. https://doi.org/10.1093/nar/gkv1239.

75. Pal C, Bengtsson-Palme J, Rensing C, Kristiansson E, Larsson DGJ. 2014. BacMet: antibacterial biocide and metal resistance genes database. Nucleic Acids Research 42:D737–D743. https://doi.org/10.1093/nar/gkt1252.

76. Moriya Y, Itoh M, Okuda S, Yoshizawa AC, Kanehisa M. 2007. KAAS: an automatic genome annotation and pathway reconstruction server. Nucleic Acids Research 35:W182–W185. https://doi.org/10.1093/nar/gkm321.

77. Carbon S, Dietze H, Lewis SE, Mungall CJ, Munoz-Torres MC, Basu S, Chisholm RL, Dodson RJ, Fey P, Thomas PD, Mi H, Muruganujan A, Huang X, Poudel S, Hu JC, Aleksander SA, McIntosh BK, Renfro DP, Siegele DA, Antonazzo G, Attrill H, Brown NH, Marygold SJ, McQuilton P, Ponting L, Millburn GH, Rey AJ, Stefancsik R, Tweedie S, Falls K, Schroeder AJ, Courtot M, Osumi-Sutherland D, Parkinson H, Roncaglia P, Lovering RC, Foulger RE, Huntley RP, Denny P, Campbell NH, Kramarz B, Patel S, Buxton JL, Umrao Z, Deng AT, Alrohaif H, Mitchell K, Ratnaraj F, Omer W, Rodriguez-Lopez M, et al. 2017. Expansion of the Gene Ontology knowledgebase and resources. Nucleic Acids Research 45:D331–D338. https://doi.org/10.1093/nar/gkw1108.

78. Carbon S, Douglass E, Dunn N, Good B, Harris NL, Lewis SE, Mungall CJ, Basu S, Chisholm RL, Dodson RJ, Hartline E, Fey P, Thomas PD, Albou LP, Ebert D, Kesling MJ, Mi H, Muruganujian A, Huang X, Poudel S, Mushayahama T, Hu JC, LaBonte SA, Siegele DA, Antonazzo G, Attrill H, Brown NH, Fexova S, Garapati P, Jones TEM, Marygold SJ, Millburn GH, Rey AJ, Trovisco V, dos Santos G, Emmert DB, Falls K, Zhou P, Goodman JL, Strelets VB, Thurmond J, Courtot M, Osumi-Sutherland D, Parkinson H, Roncaglia P, Acencio ML, Kuiper M, Laegreid A, Logie C, Lovering RC, et al. 2019. The Gene Ontology Resource: 20 years and still GOing strong. Nucleic Acids Research 47:D330-D338. https://doi.org/10.1093/nar/gky1055.

79. Huerta-Cepas J, Forslund K, Coelho LP, Szklarczyk D, Jensen LJ, von Mering C, Bork P. 2017. Fast Genome-Wide Functional Annotation through Orthology Assignment by eggNOG-Mapper. Molecular Biology and Evolution 34:2115–2122. https://doi.org/10.1093/molbev/msx148.

80. Huerta-Cepas J, Szklarczyk D, Heller D, Hernandez-Plaza A, Forslund SK, Cook H, Mende DR, Letunic I, Rattei T, Jensen LJ, von Mering C, Bork P. 2019. eggNOG 5.0: a hierarchical, functionally and phylogenetically annotated orthology resource based on 5090 organisms and 2502 viruses. Nucleic Acids Research 47:D309–D314. https://doi.org/10.1093/nar/gky1085.

81. Ye J, Zhang Y, Cui HH, Liu JW, Wu YQ, Cheng Y, Xu HX, Huang XX, Li ST, Zhou A, Zhang XQ, Bolund L, Chen Q, Wang J, Yang HM, Fang L, Shi CM. 2018. WEGO 2.0: a web tool for analyzing and plotting GO annotations, 2018 update. Nucleic Acids Research 46:W71-W75. https://doi.org/10.1093/nar/gky400.

82. Kang DWD, Li F, Kirton E, Thomas A, Egan R, An H, Wang Z. 2019. MetaBAT 2: an adaptive binning algorithm for robust and efficient genome reconstruction from metagenome assemblies. Peerj 7:e7359. https://doi.org/10.7717/peerj.7359.

83. Wu YW, Simmons BA, Singer SW. 2016. MaxBin 2.0: an automated binning algorithm to recover genomes from multiple metagenomic datasets. Bioinformatics 32:605–607. https://doi.org/10.1093/bioinformatics/btv638.

84. Alneberg J, Bjarnason BS, de Bruijn I, Schirmer M, Quick J, Ijaz UZ, Lahti L, Loman NJ, Andersson AF, Quince C. 2014. Binning metagenomic contigs by coverage and composition. Nature Methods 11:1144–1146. https://doi.org/10.1038/nmeth.3103.

85. Parks DH, Imelfort M, Skennerton CT, Hugenholtz P, Tyson GW. 2015. CheckM: assessing the quality of microbial genomes recovered from isolates, single cells, and metagenomes. Genome Research 25:1043–1055. https://doi.org/10.1101/gr.186072.114.

86. Creevey CJ, Doerks T, Fitzpatrick DA, Raes J, Bork P. 2011. Universally Distributed Single-Copy Genes Indicate a Constant Rate of Horizontal Transfer. Plos One 6:e22099. https://doi.org/10.1371/journal.pone.0022099.

87. Jain C, Rodriguez-R LM, Phillippy AM, Konstantinidis KT, Aluru S. 2018. High throughput ANI analysis of 90K prokaryotic genomes reveals clear species boundaries. Nature Communications 9:5114. https://doi.org/10.1038/s41467-018-07641-9.

88. Price MN, Dehal PS, Arkin AP. 2010. FastTree 2-Approximately Maximum-Likelihood Trees for Large Alignments. Plos One 5:e9490. https://doi.org/10.1371/journal.pone.0009490.

89. Letunic I, Bork P. 2019. Interactive Tree Of Life (iTOL) v4: recent updates and new developments. Nucleic Acids Research 47:W256–W259. https://doi.org/10.1093/nar/gkz239.

90. Yin YB, Mao XZ, Yang JC, Chen X, Mao FL, Xu Y. 2012. dbCAN: a web resource for automated carbohydrate-active enzyme annotation. Nucleic Acids Research 40:W445–W451. https://doi.org/10.1093/nar/gks479.

91. Lannes R, Olsson-Francis K, Lopez P, Bapteste E. 2019. Carbon Fixation by Marine Ultrasmall Prokaryotes. Genome Biology and Evolution 11:1166–1177. https://doi.org/10.1093/gbe/evz050.

92. Blin K, Shaw S, Steinke K, Villebro R, Ziemert N, Lee SY, Medema MH, Weber T. 2019. antiSMASH 5.0: updates to the secondary metabolite genome mining pipeline. Nucleic Acids Research 47:W81–W87. https://doi.org/10.1093/nar/gkz310.

93. Edgar RC. 2004. MUSCLE: multiple sequence alignment with high accuracy and high throughput. Nucleic Acids Research 32:1792–1797. https://doi.org/10.1093/nar/gkh340.

